# Development of a Novel Peptide Aptamer that Interacts with the eIF4E Capped-mRNA Binding Site using Peptide Epitope Linker Evolution (PELE)

**DOI:** 10.1101/2022.02.17.480295

**Authors:** Yuri Frosi, Simon Ng, Yen-Chu Lin, Shimin Jiang, Siti Radhiah Ramlan, Dilraj Lama, Chandra S Verma, Ignacio Asial, Christopher J Brown

**Affiliations:** Disease Intervention Technology Lab (DITL), IMCB (A*STAR), 8A Biomedical Grove, #06-04/05, Neuros/Immunos, Singapore 138648; Insilico Medicine Taiwan Ltd., Suite 2013, No.333, Sec.1, Keelung Rd., Xinyi Dist. 110, Taipei, Taiwan; Department of Microbiology, Tumor and Cell Biology, Karolinska Institutet, Biomedicum Quarter 7B-C Solnavägen 9, 17165 Solna, Sweden; Bioinformatics Institute (A*STAR), 30 Biopolis Street, #07-01 Matrix, Singapore 138671; DotBio, 1 Research Link, Singapore 117604; Nanyang Technological University, School of Biological Sciences

## Abstract

Identifying new binding sites and poses that modify biological function are an important step towards drug discovery. We have identified a novel disulphide constrained peptide that interacts with the cap-binding site of eIF4E, an attractive therapeutic target that is commonly overexpressed in many cancers and plays a significant role in initiating a cancer specific protein synthesis program though binding the 5’cap (7’methyl-guanoisine) moiety found on mammalian mRNAs. The use of disulphide constrained peptides to explore intracellular biological targets is limited by their lack of cell permeability and the instability of the disulphide bond in the reducing environment of the cell, loss of which results in abrogation of binding. To overcome these challenges, the cap-binding site interaction motif was placed in a hypervariable loop on an VH domain, and then selections performed to select a molecule that could recapitulate the interaction of the peptide with the target of interest in a process termed Peptide Epitope Linker Evolution (PELE). A novel VH domain was identified that interacted with the eIF4E cap binding site with a nanomolar affinity and that could be intracellularly expressed in mammalian cells. Additionally, it was demonstrated to specifically modulate eIF4E function by decreasing cap-dependent translation and cyclin D1 expression, common effects of eIF4F complex disruption.

## Introduction

Uncovering novel binding sites and poses that modulate biological function is a defining goal of lead discovery to facilitate therapeutic development. The molecular characteristics of protein surfaces, ranging from planar interactions surfaces, to highly dynamic and plastic surfaces, can pose significant challenges to small molecule discovery ^1–4^. Peptides are ideal modalities for identifying new binding sites due to their ability to adopt multiple configurations, mimic the molecular features found at protein binding interfaces, and to interact with their target molecules with relatively high affinities and specificities.^5,6^ The range of potential binding sites open to peptide binding can be extended further by constraining their secondary structure through cyclization, this enables a reduction in the entropic cost of binding for interactions modes poorly sampled by linear peptide ensembles.

The translation of peptide compounds into suitable molecules for lead development and biological evaluation face several hurdles: 1) limited intracellular permeability and 2) proteolytic instability.^1,5^ The process to surmount this challenge is highly resource- and time-consuming. Therefore, to understand the biological and phenotypical consequences of the binding modes of these molecules further in a reasonable timeframe, a methodology is required to present these binding epitopes in a relevant manner to enable these studies. One attractive approach is to embed these binding epitopes within the context of a small and highly stable protein backbone (termed scaffold), where they will be doubly constrained at their N and C terminals. These modalities are usually referred to as peptide aptamers (PA).^7^ An advantageous property of PAs are that they are genetically encodable and can be expressed within mammalian cell lines and animal models to facilitate advanced studies.^7,8^ PAs are usually designed via 2 distinctive approaches either by insertion of a single sequence into a loop region (“loop on frame”) or via mutations of specific residues embedded in rigid secondary structural (“rigid motif) elements within the selected scaffold protein. ^7,8^

These approaches in addition to rational protein engineering techniques are both amenable to combinatorial display methods such as yeast and phage display. However, they have limitations: 1) specific or hypervariable loop sequences are inserted into scaffolds that are not optimized to stabilise their bound conformations, whilst 2) the use of “rigid motifs” leads to sampling limited areas of 3-dimensional space e.g., mutations that lie down one side of an α-helix. To circumvent these issues a 2-step discovery process (termed **<u>P</u>**eptide **<u>E</u>**pitope **<u>L</u>**inker **<u>E</u>**volution, PELE) was utilized (**Figure 1A**), whereby the surface of the target protein is mapped for novel peptides binders utilizing constrained and linear peptide phage libraries, where upon any isolated interaction motifs are inserted into a larger hypervariable loop located on a selected scaffold protein. Libraries are then constructed with the interaction motif (‘peptide epitope’) located at different positions within the hypervariable loop, and then the selection against the target protein re-performed to select for sequences (‘linker evolution’) that optimally present the interaction motif within the context of the scaffold. The advantage of this approach over generating naïve libraries is that the diversity of the libraries in both steps are now only focussed on finding the optimal solution for one variable each, ‘binding’ and ‘presentation’, respectively, rather than simultaneously in a single step. Thus, allowing for more effective searching of sequence space especially with techniques that attain lower thresholds of diversity (e.g., yeast ~ 10^6-7^ vs mRNA display ~ 10^12-13^).

**Figure 1:**
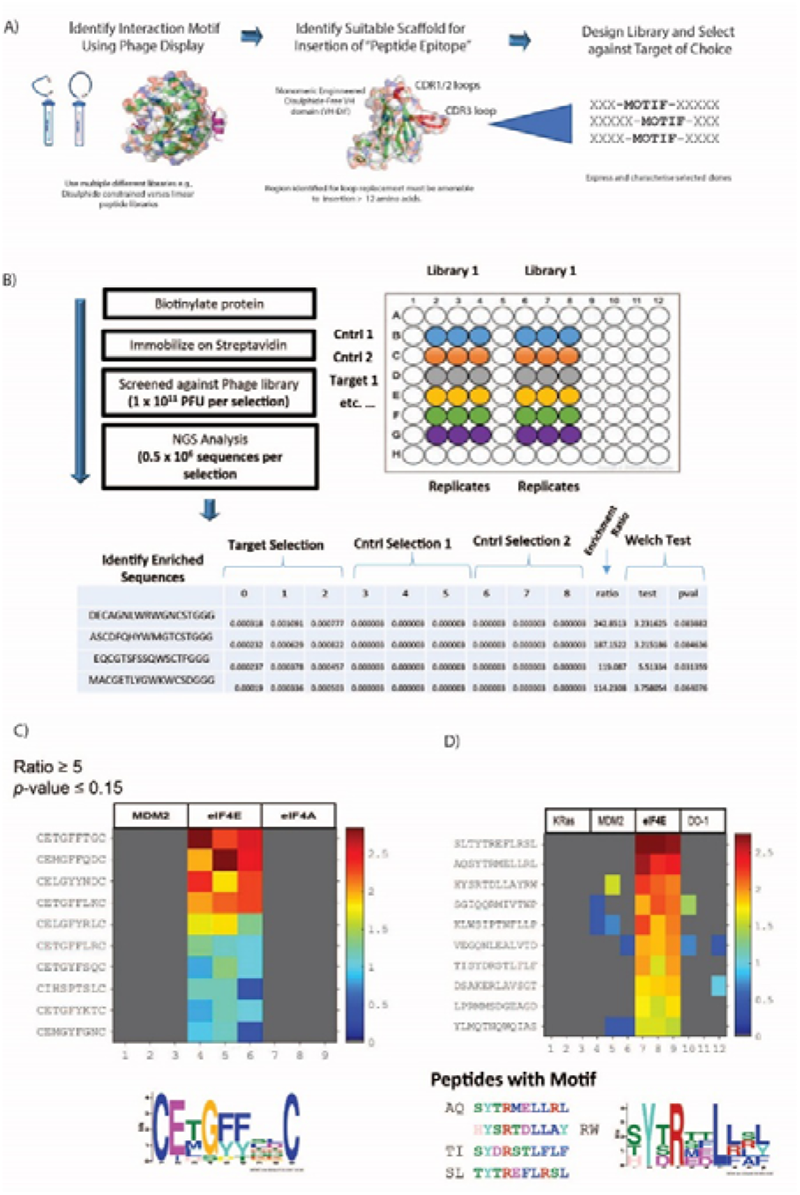
**A)** The schematic outlines the process termed **<u>P</u>**eptide **<u>E</u>**pitope **<u>L</u>**inker **<u>E</u>**volution (PELE). Peptide phage display libraries are used to probe the surface of the target protein to discover new binding motifs and modalities (e.g., linear or constrained libraries). Upon identification, the novel motif or modality can be inserted into a larger hypervariable loop located on a selected scaffold protein Several distinctive libraries can then constructed be with the interaction motif (‘**p**eptide **e**pitope’) located at different positions within the hypervariable loop, and then the selection against the target protein re-performed to select for sequences (‘**l**inker **e**volution’) that optimally present the interaction motif within the context of the scaffold. **B)** A brief outline of next generation sequencing (NGS) enhanced phage display. A selected phage library is panned against an immobilised target protein using three technical repeats and in parallel against corresponding negative control selections where the target protein is either removed or replaced with a different target protein. Bound phage is then eluted, amplified and sequenced using NGS protocols (NextSEQ, Illumina). The FASTQ file generated from the sequencing data was processed by in-house PYTHON scripts that identified the barcodes and constant flanking regions and extracted the reads of the correct length corresponding to the variable peptide library. The table presents the list of sequences identified from each selection with their associated abundance. The abundance is calculated by taking the copy number of each sequence and normalizing it by dividing the copy number by the total number of reads in each sequence. Sequences not observed in a specific replicate were assigned a copy number of zero. The enrichment ratio of each sequence in the target selection was calculated by determining the mean fraction from the target screen replicates and dividing it by the mean fraction from the selected control screen replicates. Since the denominator must not be a zero when taking the ratio, sequences with zero copy number found in all three replicates are assigned with an arbitrary copy number before taking the normalization. Significance of the ratio was assessed using one-tailed, unequal variance Welch test. A heat map (**C and D**) is then generated to identify the enriched peptides that have ratio and p-values corresponding to the parameters stated in the figure. Each individual colour coded block on the map represents the abundance of the unique sequence in each selection and the sequence are ordered by their ratio value. **C)** Heatmap showing sequences enriched from the M13 disulphide constrained 7mer library (C7C) against eIF4E, but not in the 2 control selections (Mdm2 and eIF4A). **D)** Heatmap showing sequences enriched from the M13 linear 12mer library against eIF4E, but not in the 3 control selections (Mdm2, K-RAS and DO-1). The sequence motif in **C)** was generated from the enriched sequences using MEME, whilst in **D)** it was generated from the sequences exhibiting the known eIF4E binding motif (YXXXL, X is any amino acid).

The PELE process was used to identify novel cyclic peptides that interact with the elongation initiation factor 4E (eIF4E) at the capped-mRNA binding site and to evolve the identified interaction motif onto an autonomous, disulphide-free VH domain.^9^ eIF4E itself plays a critical role in cap-dependent translation and is an attractive target for anti-oncogenic therapy as it is frequently overexpressed and/or mis-regulated in many tumours, is associated with poor prognosis in patients, is implicated in chemoresistance^10–12^ and also orchestrates a cancer specific mRNA translational program e.g., c-myc, cyclin-D1.^13,14^ eIF4E binds the cap structure found at the 5’ end of mammalian mRNA as part of the eIF4F complex, whereupon the complex interacts with the initiation factor eIF3 via the eIF4G subunit, which enables the recruitment of the 43S pre-initiation complex (PIC). The PIC complex then transverses the 5⍰ UTR of the mRNA until it encounters the AUG start codon, where it initiates protein synthesis though formation of the 80s ribosomal complex.^15^ The formation of the eIF4F complex in normal cells is finely controlled by the 4E-BP proteins (4EBP1, 4EBP2 and 4EBP3), where they competitively inhibit the interaction of eIF4E with eIF4G. Both the 4E-BPs and eIF4G1 share a common eIF4E interaction motif (YXXXXLΦ, X = any amino acid and Φ = hydrophobic amino acid), which interacts at the same site on eIF4E.^16^ Disruption of eIF4G binding impairs the assembly of the eIF4F complex on the 5⍰ cap structure and prevents cap-dependent translation. The actions of 4EBPs are controlled by their phosphorylation status, which is under the specific control of mTOR.^17^ Aberrant eIF4E activity in cancer can be targeted using several approaches. One method is to disrupt eIF4F formation using inhibitors of the PI3K/AKT/mTORC pathway ^18^, which induce de-phosphorylation of the 4EBPs^15^. Another strategy is to prevent eIF4E phosphorylation through the use of MNK1/2 kinase antagonists in the RAS/ERK signalling pathway ^15^. Other approaches that have been used to antagonise the activity of the eIF4F complex are ASOs (antisense oligonucleotides) to reduce eIF4E expression, inhibitors of the eIF4A helicase^15^ and development of cap analogues to prevent binding of capped mRNA.^15,19,20^ Additionally, small molecules and peptides can be used to target the interface between eIF4G and eIF4E.^15,21,22^

In this manuscript, we report the first peptide inhibitor of the eIF4E cap binding site and have used this to develop a novel peptide aptamer suitable for intracellular expression and therapeutic modelling studies. The structures of both the peptide and the peptide aptamer (PA) with eIF4E revealed a novel interaction with the cap-binding site of eIF4E that provides a unique template for future small molecule and peptidomimetic design strategies. Additionally, the VH domain-based PA was also used to construct a unique split luciferase with the potential to screen eIF4E cap antagonists directly in live cells and support future cap-analogue mimetic discovery.

## Results and Discussion

### Discovery of a Novel eIF4E Cap-Binding Cyclic Peptide Binding Motif

Two M13 phage peptide libraries (New England Biolabs) either consisting of 7 randomised amino acids constrained by a disulphide bond formed between 2 cysteine residues (ACXXXXXXXC, X = any amino) or a hypervariable linear 12mer sequence were panned against biotinylated eIF4E. Parallel selections were performed concurrently against several negative control biotinylated proteins also. The panning culminated in single round to avoid selection of fast-growing phage clones.^23^ Recovered phage populations were then subjected to NGS (Next Generation Sequencing, Illumina NextSEQ technology) sequencing, whereupon differential enrichment analysis was performed to identify peptide sequences that specifically bound eIF4E over the control proteins (**figure 1B**). The 12mer library selection identified peptides with the interaction motif (YXRXXL[L/R/F])), which is highly similar to the well-known eIF4E binding motif (YXXXXLΦ, Φ is any hydrophobic amino acid).^16^ In contrast, the motif enriched in the disulphide constrained peptide selection isolated a previously unknown putative eIF4E interaction motif (CE[M/L/T]G[F/Y]XXC) (**figure 1C and 1D**).

In the absence of any sequence similarity with described eIF4E interacting proteins (eIF4G1 and the 4E-BP family) and the eIF4E interaction motif (enriched in the 12mer selection), competitive fluorescence anisotropy experiments were performed to delineate the binding sites of the EE-01 to EE-09 peptides on eIF4E utilising either a FAM labelled m^7^GTP (m^7^GTP^FAM^) or a FAM labelled eIF4G/4E-BP1 site interacting peptide (eIF4G^FAM^) (**figure 2A, 2B and 2C**). None of the disulphide constrained peptides were observed to displace eIF4G^FAM^ from eIF4E, however several cyclic peptides did compete for binding at the cap-site with m^7^GTP^FAM^. EE-02 was determined to interact with eIF4E with a K^d^ of 406.2 ± 3.6 nM, 12-fold more strongly than the next best performing cyclic peptide (EE-09, K^d^ = 4,860 ± 633.4 nM). Alanine scanning mutagenesis experiments confirmed that the residues conserved in the disulphide constrained peptide motif (C^2^E^3^[M/L/T]^4^G^5^[F/Y]^6^[F/Y]^7^X^8^X^9^C, **figure 1A**) were all necessary for binding (**table 1**). Mutation of Q8A in the EE-02 sequence had no effect on binding to eIF4E, whilst the replacement of D9A resulted in a small attenuation of the K_d_ (**table 1**). Additionally, the dependence of EE-02 binding with eIF4E upon the disulphide constraint was demonstrated to be critical under reducing conditions.

**Figure 2:**
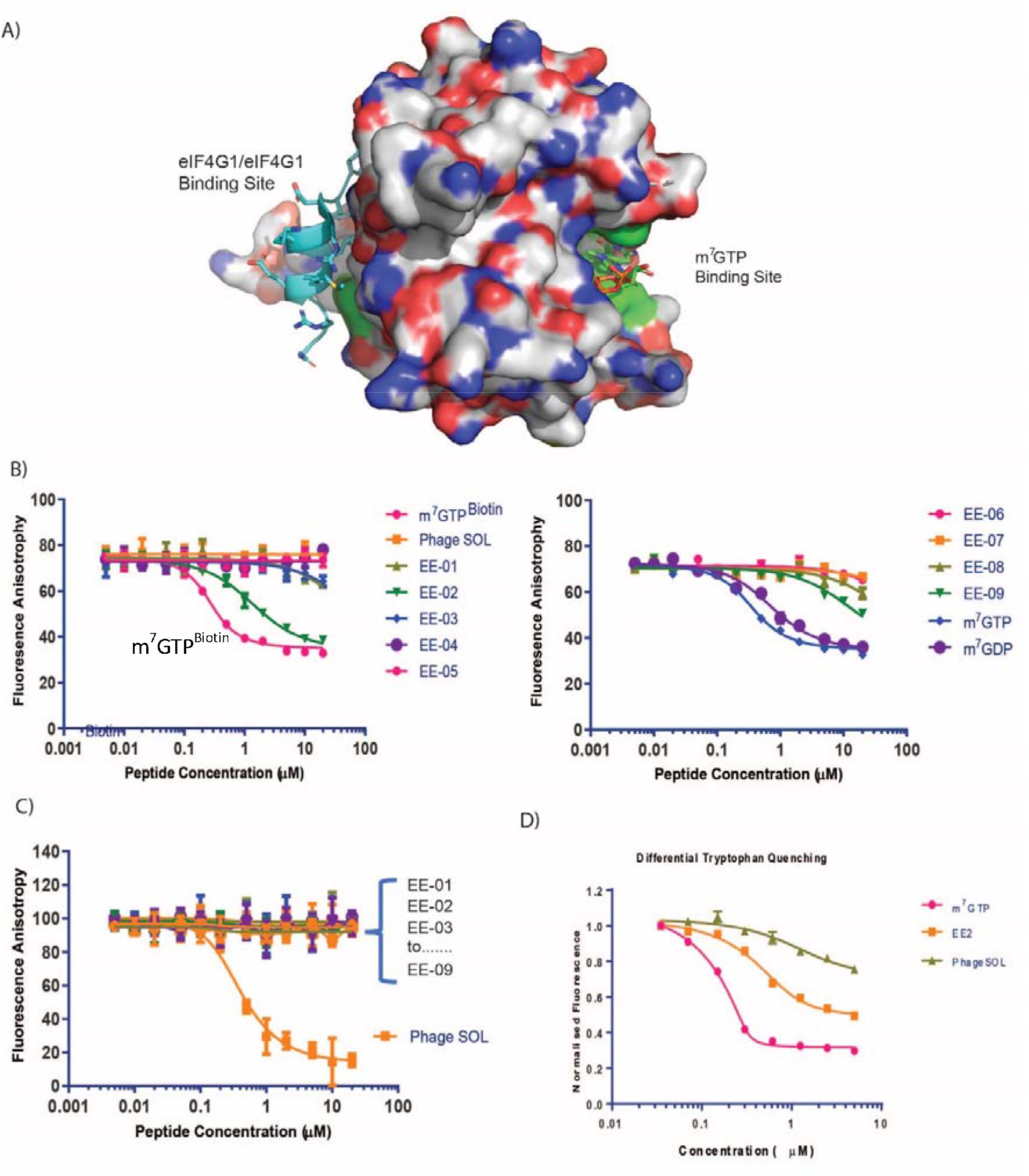
**A)** A surface representation of eIF4E depicting the location of the m^7^GTP (capped mRNA) and eIF4G1/4E-BP1 binding sites. Locations of tryptophan residues whose intrinsic fluorescence is sensitive to binding by either m^7^GTP or peptides that interact with the eIF4G binding site are shown in green. **B)** Competitive fluorescence anisotropy experiments with FAM labelled m^7^GTP assessing binding of the cyclic peptides to the cap-binding site. **C)** Competitive fluorescence anisotropy experiments with FAM labelled eIF4G1 derived peptide assessing binding of the cyclic peptides to the eIF4G/4E-BP1 binding site. Apparent K_d_s (**see table 1**) were determined by curve-fitting using Prism (Graphpad, LtD). See materials and methods. **D)** eIF4E intrinsic tryptophan fluorescence was assessed in response to titrations of m^7^GTP, PHAGESOL (Ac-KKRYSR*QLL*-NH_2_) and EE-02, respectively.

**Table 1:**
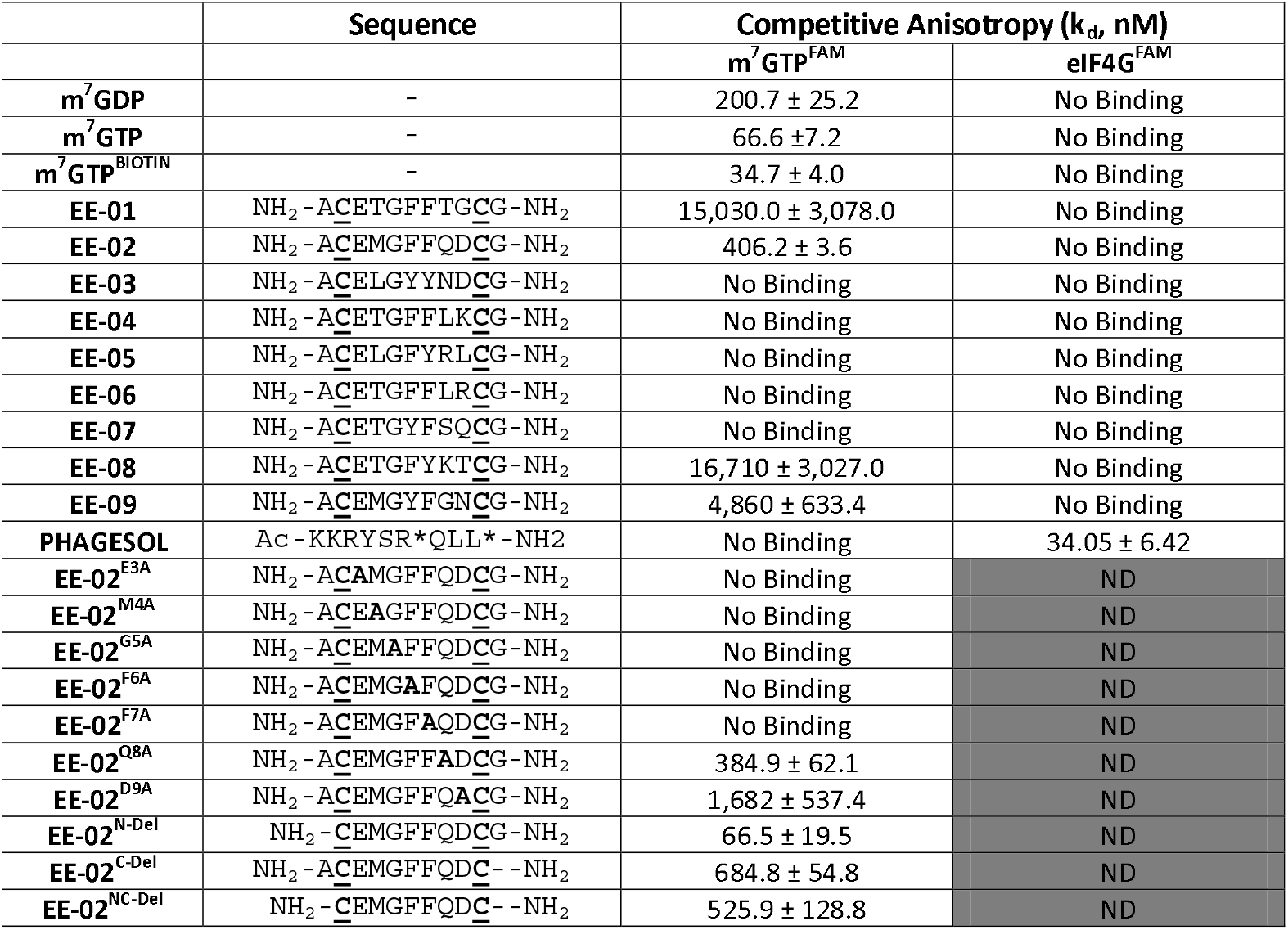
Binding assessment of disulphide constrained peptides isolated using M13 phage display against eIF4E (EE-02 to EE-09) and alanine scanning mutants exploring the interaction profile of EE-02 under non-reducing conditions. The binding sites of the peptides EE-01 to EE-09 were mapped onto eIF4E using two competitive based fluorescence anisotropy assays, one of which used a FAM labelled m^7^GTP (m^7^GTP^FAM^) to monitor for binding at the cap-binding site, whilst the other assay used a FAM labelled canonical site interacting peptide (eIF4G^FAM^) to measure binding at the eIF4G interaction site. Dissociation constants were determined using a 1:1 binding model and are described in the materials and methods. m^7^GTP, m^7^GDP and m^7^GTP^BIOTIN^ were used as positive controls for the m7GTP^FAM^ competition assay, whilst PHAGESOL was used as a positive control for the eIF4G^FAM^ competition assay. EE-02 alanine mutant derivatives were only assessed for binding in the m^7^GTP^FAM^ competition assay. ND = Not determined. K_d_s > 20,000 were denoted as non-binders. Experiments were performed in triplicate.

Tryptophan quenching experiments were performed to compare the binding mechanisms of EE-02 with m^7^GTP and PHAGESOL (a phage modified eIF4E interacting peptide that binds at the eIF4G binding site)^24,25^ against eIF4E. It is well established that the binding of m^7^GTP to eIF4E results in significant tryptophan fluorescent quenching of W56 and W102 (**figure 2A**), both involved in recognising the m^7^G moiety of m^7^GTP (**figure 2D**), whilst peptides that interact with eIF4E at the eIF4G site also result in quenching of W73 (**figure 2A and 2D**).^26^ However, the reduction in total eIF4E fluorescence caused by peptide binding is significantly less compared to m^7^GTP binding at the cap-site (**figure 2D**). In contrast, the quenching of intrinsic tryptophan fluorescence by EE-02 produces a significantly different profile to both m^7^GTP and PHAGESOL (**figure 2D**), indicating that EE-02 binds via a mechanism substantially different to both.

### The Constrained Macrocyclic Peptide EE-02 Interacts with eIF4E Cap-Binding Site via a Unique Binding Pose

The structure of the EE-02 complex was solved using X-ray crystallography confirming EE-02 bound to eIF4E at the cap-binding site (**figure 3A**), but more interestingly revealed that the site had undergone substantial conformational changes compared to the structure of cap-bound eIF4E (**figure 3B**).^27,28^ These changes were primarily localised to the W56 containing loop (48-60) with smaller sidechain structural changes occurring elsewhere around the pocket. The net effect of these changes were that the side chains of W56 and W102 that play critical roles in recognising the m^7^G moiety of the cap-analogue no longer reside within the cap-binding site and make contrasting interactions with EE-02 compared to m^7^GTP. The differences between the EE-02:eIF4E and m^7^GTP:eIF4E complex structures also explain the substantial differences observed in their tryptophan quenching profiles (**figure 2D**).

**Figure 3:**
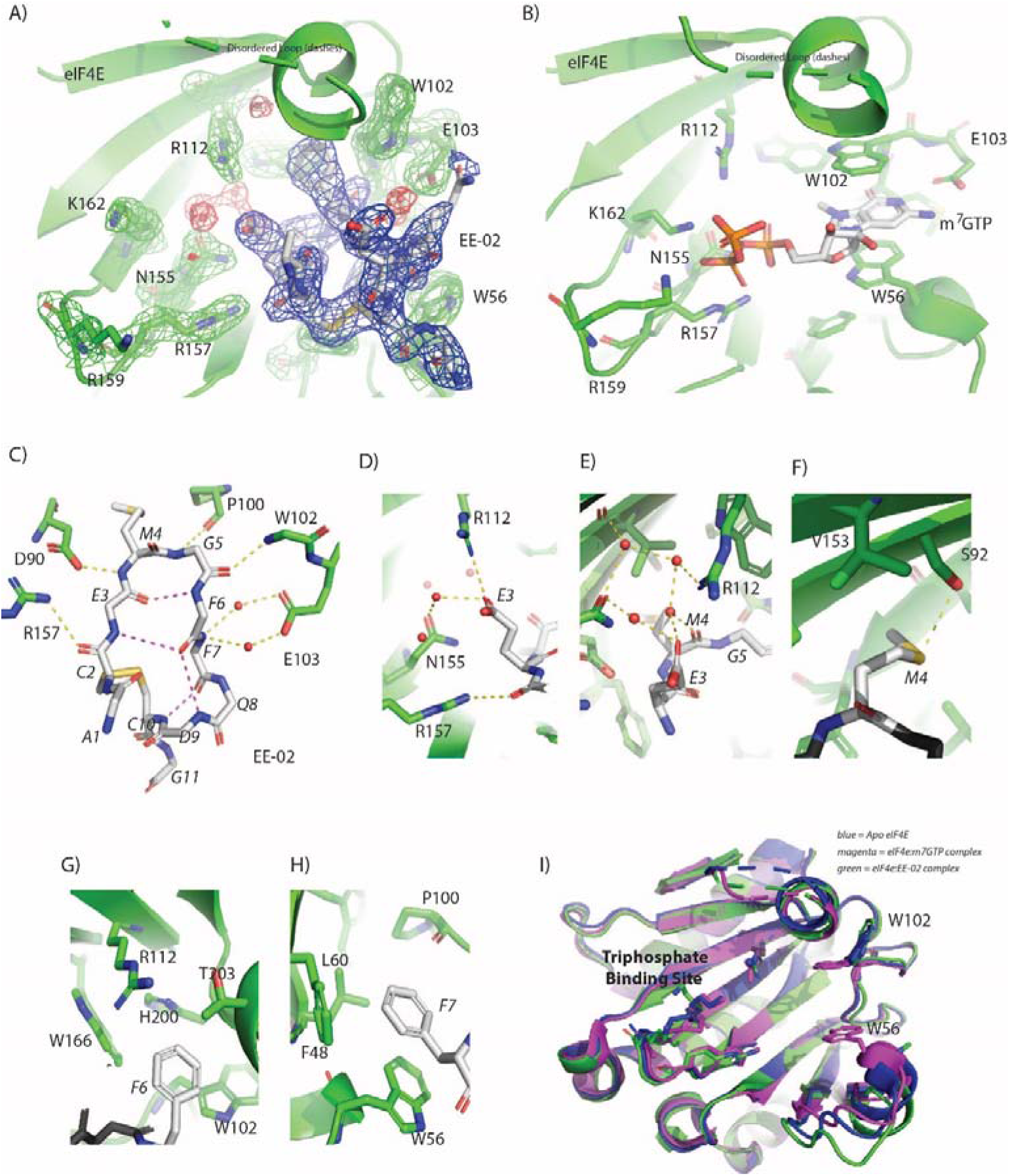
**A)** The 2F_o_-F_c_ electron density map (1.2σ) showing the EE-02 disulfide constrained peptide bound to eIF4E at the cap-binding site. EE-02 is highlighted in the blue mesh, structured waters in the red mesh and eIF4E interacting residues with green mesh. **B)** Complex of eIF4E bound to m^7^GTP (PDB ID: 2V8W) indicating conformational differences with the EE-02:eIF4E complex structure. **C)** EE-02 when bound to eIF4E forms a β-hairpin turn-like structure that is stabilized by intra hydrogen bonds between the backbone carbonyl of E3 and backbone amide of F6, and the backbone amide of E-03 and carbonyl of F6 (3.1⍰ and 4.1⍰, respectively). The conformation of the cyclic peptide is further rigidified by hydrogen bonds between the C10 amide and the carbonyl of F7, and the backbone of N9 and the carbonyl of F7 (3.0⍰ and 3.8⍰, respectively). The polypeptide backbone of EE-02 also forms a set of critical interactions with eIF4E (< 3.2⍰) shown in yellow. **D)** to **G)** show the interactions made by the conserved residues of the cyclic peptide interaction motif in EE-02 (white) with eIF4E (green). **D)** E3 electrostatically interacts with R112 and forms a water mediated hydrogen bond interaction with N155. **E)** The carbonyl group of G5 forms no direct interactions with eIF4E but forms a hydrogen bond with a structured water, which is part of a larger network of structured waters that facilitates the interaction of EE-02 with eIF4E. **F)** M4 forms a dipole interaction with the hydroxyl group of S92 and a variety of hydrophobic contacts with residues F48, W46, L60 and P100. **G)** F6 forms hydrophobic contacts with the residues T203 A204, H200, W166 and W102 of eIF4E. **H)** stacking interactions with W56 and edge on face interactions with F48. Additionally, it forms a hydrophobic contact with P100. **I)** Overlay of the EE-02:eIF4E complex with unbound eIF4E (PDB ID: 4BEA) and m7GTP bound eIF4E demonstrating the similarity of the EE-02 bound conformation to the apo structure. Ligands interacting at the cap binding site (EE-02 and m GTP are not shown for clarity).

The EE-02 peptide forms a β-hairpin turn-like structure in the binding pocket that allows the side chains of the constrained peptide motif to efficiently interact with eIF4E (**figure 3C**). The glycine at position 5 due to its steric permissiveness enables optimal formation of the β-turn type structure, and in turn a stabilising intramolecular h-bond between the backbone carbonyl of E3 and backbone amide of F6. The E3 of the selected motif (E^3^MGFF^7^) forms direct electrostatic interactions with R112 of eIF4E (figure 3D), whilst M4 forms hydrophobic contacts with the back of the binding pocket and a specific hydrogen bond with S92 via its sulphur atom (**figure 3E**). Residue F6 forms a range of hydrophobic interactions (3.6Å - 4.2Å) that include T203 A204, H200, W166 and W102 of eIF4E (**figure 3F**). In contrast, F7 forms stacking interactions with W56 and edge on face interactions with F48 (**figure 3G**). Additional main-chain interactions are also formed by EE-02 that contribute to the energetics of binding with eIF4E: the backbone carbonyl of C2 interacts with the R157 sidechain (**figure 3C**), the carbonyl and amide back bones of G5 and F7, respectively, form water mediated interactions with the carboxylic group of E103 (**figure 3C**), and the backbone carbonyl of M4 interacts with a structure water network that involves h-bonds with N155 and R112 (**figure 3G**). Binding energy decomposition analysis from MD simulations of the eIF4E:EE-02 complex structure demonstrates that M4, F6 and F7 contribute a significant proportion of the binding energy (**Figure S1A**). The part of the cap-binding that recognizes the triphosphate tail (R157, K159 and K162) of m^7^GTP is not involved in binding EE-02, and undergoes negligible structural changes. Interestingly, the conformation of the eIF4E cap-binding site when bound to EE-02 is very similar to its unbound configuration^21,29,30^, where the W102 side chain and the W56 loop also rotate and swing out of the cap-binding site (**figure 3I**). MD simulations of the unbound EE-02 were performed, indicating that its structure does not vary dramatically from the bound form and overall retains a similar fold to that observed in the crystal form (**figure S2A and S2B**).

### Design and Development of a Novel Miniprotein that Interacts with the Cap-Binding Site of eIF4E

The use of disulphide constrained peptides to evaluate intracellular biological targets is severely limited by their inherent lack of cell permeability and the instability of the disulphide bond in the reducing environment of the cell. To overcome these limitations, the EE-02 binding epitope was grafted into the CDR3 loop region of an engineered monomeric VH-domain, termed DiF-VH.^9^ The DiF-VH scaffold has several attractive features: 1) relatively large peptide insertions can be made into the CDR3 loop region and 2) the protein scaffold is amenable to expression in E.Coli and mammalian cells,. Additionally, the points where the CDR3 loop initiates and terminates itself in the VH domain are spatially close together, suggesting that the protein scaffold can act as structural constraint that mimics the function of the disulphide bond in the cyclic peptide (**figure 4A**).

**Figure 4:**
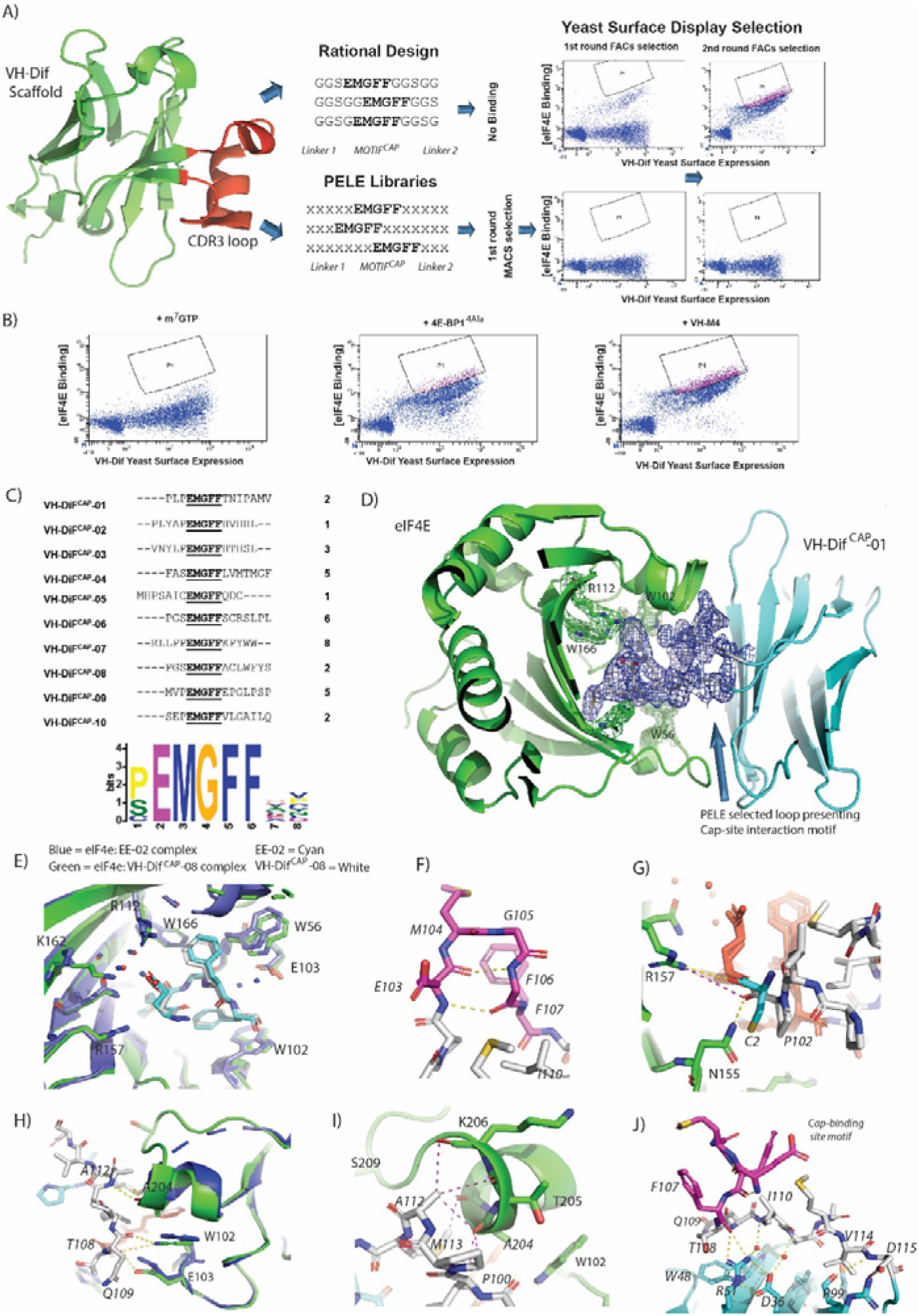
**A)** The CDR3 region (highlighted in red) of the VH-DiF scaffold (PDB ID: 7D8B) was selected for replacement by rationally designed loops. The engineered loops were designed to present the EE-02 motif in the correct conformation to interact with eIF4E using polyGly linkers. However, the VH-DiF derived proteins, when tested, exhibited no binding to eIF4E. Peptide Epitope Loop Exchange (PELE) libraries were also constructed and inserted at the same site in the VH-DiF scaffold. Optimal linkers needed to present the EE-02 motif correctly for binding were selected by YSD (**Y**east **S**urface **D**isplay). The YSD (yeast surface display) selection against eIF4E went through an initial round of selection performed with IMACs, followed by 2 rounds of in-solution selection using flow cytometry to enrich the population for high affinity eIF4E binders, where biotinylated eIF4E was detected using dye-labelled streptavidin. Insets show the enrichment in eIF4E cap-binders in the PELE library after rounds 2 and 3 of FACs selection. Negative control experiments were performed with the same library inputs that showed negligible non-specific binding within the enriched populations in the absence of eIF4E. **B)** Samples from the final round input for YSD selection were co-incubated with either m^7^GTP, 4E-BP1^4ALA^ or VH-M4 in order to compete with the VH-DiF population enriched for eIF4E binding with biotinylated eIF4E (measured in **A**). A significant reduction in the enriched population interacting with eIF4E only occurred with m^7^GTP treatment indicating that the selected eIF4E binders were specific for the cap-binding site. **C)** The table lists the 10 unique VH-DiF sequences identified from the 34 yeast clones sequenced in the final round of YSD selection, with their corresponding frequencies. A recognition motif was generated from the identified sequences using MEME (XXX), which in addition to showing the invariant cyclic peptide interaction motif, also identified that proline was preferentially enriched for at the position immediately preceding the motif. **D)** Complex structure of eIF4E (green) with VH-DiF^CAP^-01 (cyan) highlighting the binding of the PELE selected motif presenting linkers to eIF4E. The 2F_o_-F_c_ electron density map of the cap interacting loop structure is shown in blue (1.2σ). **E)** Overlay of the cap binding motif of VH-DiF^CAP^-01 (E^103^MGFF^107^, shown in cyan) with the equivalent residues in EE-02 (shown in white) highlighting the loss of the water mediated interactions between EE-02 and E-103 (blue) and a small conformation change in E103, where the interaction with R112 and the structured water network are retained. However, it does result in an additional interaction with K162 not observed in the eIF4E:EE-02 complex. **F)** The cap binding motif of VH-DiF^CAP^-01 (E^103^MGFF^107^,) forms a similar β-hairpin-like structure to that seen in the eIF4E:EE-02 complex. Additionally, the two intra backbone hydrogen bonds that formed to stabilize the bound structure of the EE-02 cyclic peptide (figure 3C) are also observed in the VH-DiF^CAP^-01 complex with eIF4E. **G)** The hydrogen bond formed between C01 of EE-02 (cyan) with R157 is not observed in the eIF4E: VH-DiF^CAP^-01 complex, where it is replaced with a hydrogen bond between P102 (white) of the ^100^PLP^102^ linker and N155. **H)** The PELE selected linker (T^108^NIPAMV^114^) form 3 hydrogen bonds with eIF4E: Residues T108 and Q109 form 2 hydrogen bonds with the indole group of W102 of eIF4E (3.7 ⍰ and 3.1 ⍰, respectively), and a hydrogen bond forms between the amide and carbonyl groups of A112 and eIF4E’s A204, respectively. **I)** Residues I110 and A113 of the linker region (T^108^NIPAMV^114^) form multiples hydrophobic contacts with residues T203, A204, T206 and G208 in eIF4E, which stabilize the α-helical secondary structure of the eIF4E region 201 to 205. J) The conformation of E^103^MGFF^107^ (highlighted red) is stabilized by a hydrophobic cluster principally formed by I110 from the linker region (T^108^NIPAMV^114^) and a salt bridge between VH-DiF^CAP^-01 residues R51 and D36, which also interact with two buried structured waters. The water network in conjunction with R51 form hydrogen bonds with the polypeptide backbone of the PELE selected loop, helping to stabilize the conformation of the cap-site interaction motif for eIF4E binding.

Initial designs where the cyclic eIF4E interaction motif (EMGFF) was introduced at different positions within a 15mer loop in the CDR3 region of the VH-DiF scaffold were tested and none exhibited any binding to eIF4E (**Figure 4A and S3**). It was then hypothesized that the use of poly-glycine linkers for epitope presentation was too permissive and did not restrain the motif sufficiently in the correct structural conformation to be able to interact with eIF4E. To overcome this issue an alternative approach termed Peptide Epitope Linker Evolution (PELE) was adopted, wherein the linker regions were randomized, and a yeast surface display (**YSD**) library generated to select for linkers that optimally displayed the eIF4E interaction motif (**Figure 4A**). To confirm that the PELE selection was successful, input samples of the enriched yeast library from the final selection round were used in competition experiments with either m^7^GTP, 4E-BP1 or VH-M4^9^ (a VH domain that binds at the eIF4G interaction site of eIF4E) against biotinylated eIF4E, which confirmed that the binding of the selectants to eIF4E were specific to the cap-binding site (**figure 4B**). 34 yeast clones from the final round of YSD were sequenced, which yielded 10 unique VH-DiF peptide aptamers (termed VH-DiF^CAP^) (**figure 4C**). Sequence analysis revealed that eIF4E interactors were isolated from each of the PELE libraries used in the selection and that proline was preferentially selected for at the amino acid position preceding the interaction motif (**figure 4C**). The VH-DiF^CAP^ peptide aptamers where then tested for bacterial expression, where upon those with good expression levels (VH-DiF^CAP^-01, VH-DiF^CAP^-02, VH-DiF^CAP^-06, VH-DiF^CAP^-09) were purified and screened for binding against eIF4E using the m^7^GTP^FAM^ competition assay (**figure S4, table 2**). The VH-DiF^CAP^ peptide aptamers that demonstrated binding in the competition assay including the constrained peptide EE-02, were then re-measured using ITC in direct binding titrations, which identified VH-DiF^CAP-01^ as the most potent eIF4E binder with a K_d_ of 35.3 ± 17.0 nM (**table 2**). A K_d_ approximately equivalent to that determined for the constrained peptide EE-02.

**Table 2:**
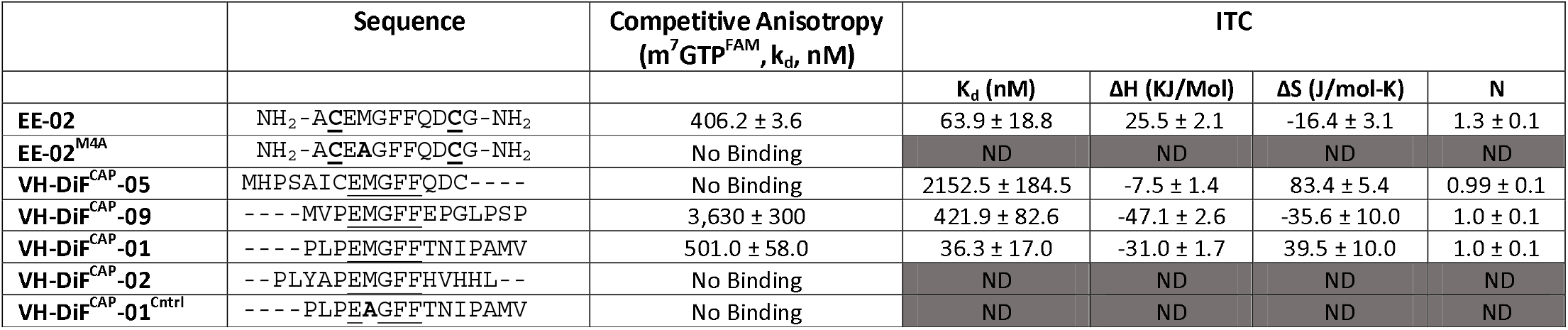
Dissociation constants were determined using both the **m**^**7**^**GTP**^**FAM**^ **competitive anisotropy assay and isothermal calorimetry (ITC) for** selected constrained peptides and peptide aptamers. ITC was also used to determine the following: enthalpy of binding (**ΔH**), entropy of (**ΔS**) binding and the stoichiometry of the interaction (N, number of binding sites). Experiments were carried out at 293 K. Experiments were performed in triplicate. ND = Not determined.

### VH-DIF ^CAP^-01 Recapitulates the Interactions of EE-02 with the Cap-Binding Site and Forms Additional Interactions

Crystallization of the VH-DIF^CAP^-01:eIF4E complex (**figure 4D**) confirmed the residues of the cyclic peptide interaction motif located in the CDR3 loop recapitulated the critical interactions observed between EE-02 and eIF4E (**figure 4E and 4F**). Additionally, binding energy binding decomposition analysis from MD simulations of the VH-DIF^CAP^-01:eIF4E complex further confirmed the similarity in the energetics of binding between EE-02 and the evolved CDR3 loop with M104, F106 and F107 again making the largest contributions to the binding energy (**Figure S1B**). The only significant deviations in the interaction of EE-02 and VH-DIF^CAP^-01 with eIF4E were: 1) a small conformational difference in the E103 sidechain and the position of its Cα backbone atom (**figure 4G**), 2) the loss of water-mediated interactions with E103 of eIF4E (**figure 4G**) and 3) a deviation in the packing of the W102 residue against F107 of VH-DIF^CAP^-01 (**figure 4G**). The re-orientation of the E103 residue is principally caused by the P^100^LP^102^ linker region of VH-DIF^CAP^-01 approaching the cap-binding site at a significantly different angle compared to the orientation of the EE-02 peptide backbone induced by the disulphide bond constraint (**figure 4F**). Interestingly, the changes observed in relation to W102 and the loss of the water-mediated interactions with E103 are primarily driven by the interactions of the evolved linker (T^108^NIPAMV^114^) with eIF4E.

The evolved linker regions of VH-DIF^CAP^-01, in addition to presenting the EE-02 interaction epitope optimally to interact with eIF4E, also forms multiple additional interactions with eIF4E (**figure 4G, 4H and 4I**). This contrasts sharply with the EE-02 cyclic peptide where only the Cys2 carbonyl group forms a hydrogen bond directly with R157 outside the residues critical for interacting with eIF4E. This this hydrogen bond does not occur in the eIF4E:VH-DiF^CAP^-01:eIF4E complex structure where but is mimicked by a hydrogen bond between the carbonyl of the P102 in the P^100^LP^102^ linker region with the side chain of N155 of eIF4e (**figure 4G**). The remainder of the N-terminal PLP linker forms no other interactions with eIF4E. However, the linker section (T^108^NIPAMV^114^) that occurs at the C-terminal of the interaction motif in VH-DIF^CAP^-01, forms most of the interactions made between eIF4E and the evolved linker regions. Residues T108 and Q109 form 2 hydrogen bonds with the indole group of W102 of eIF4E (3.7 1 and 3.1 1, respectively). An additional hydrogen bond between the linker region and eIF4E is formed between the amide and carbonyl groups of A112 and eIF4E’s A204, respectively (**figure 4H**). The remaining residues in the linker (110-114) apart from V114 make a multitude of hydrophobic contacts with residues T203, A204, T206 and G208 in eIF4E, resulting in stabilization of the α-helical secondary structure of this region of the protein (**figure 4I**). V114 in contrast is involved in interactions with the invariant part of the VH-DiF scaffold. VH-DIF^CAP^-01 also interacts weakly at a second site with eIF4E that is constituted from the CDR1 and CDR2 loops of the VH domain, where the residues S34 and S56 both form hydrogen bonds via their sidechains with the carbonyls of K52 and S53, respectively (**figure S5**).

### The Conformation of the CDR3 Interaction Loop is Stabilized by an Intricate set of Interactions

The strategy of evolving the peptide sequence regions either side of the cap binding interaction motif to enable optimal presentation of the epitope must also inherently accommodate the presence of residues that occur on the VH domain itself. A factor difficult to account for with rational design approaches. Several significant features in the linker region of the CDR3 loop that stabilized presentation of the CDR3 loop through packing interactions with the scaffold were noted: 1) the Ile110 sidechain in the linker region forms a hydrophobic core in the CDR3 loop structure that makes multiple hydrophobic contacts with other residues in the linker and the interaction motif, 2) the CDR3 loop when bound to eIF4E forms a folded structure against the R51 and D36 residue of VH-DIF^CAP^-01 which form a salt bridge between each other, 3) and that the linker residue T108 also packs directly against W48, which is found in the scaffold (**figure 4J**). The V114 residue of the evolved linker region is also involved in additional contacts with the VH domain that stabilize the fold of the CDR3 loop (**figure 4J**).

However, the most remarkable feature is the presence of two buried structure water molecules that allow the R51:D36 salt bridge to stabilize the fold of the CDR3 loop, when bound to eIF4E (**figure 4J**). These waters enable R51 and D36 to indirectly stabilize the polypeptide backbone of residue Ile110 and Q109 (**figure 4J**). Additionally, R51 also forms direct interactions with the carbonyl of F107 and the hydroxyl of the T108 sidechain, further rigidifying the presentation of the cap-site interaction motif (**figure 4J**). MD simulations also demonstrated that the unbound CDR3 loop of the VH-DiF^CAP^-01 domain undergoes a distinct structural change in comparison to the bound form (**figure S2C and S2D**). The CDR3 loop undergoes a structural relaxation, whereby the β-hairpin structure associated with the ‘EMGFF’ motif is lost and instead there is a general movement of the CDR3 loop away from the body of the VH domain scaffold (**figure 6**). Interestingly, this movement is underpinned by significant structural changes in the hydrophobic core of the CDR3 loop structure with the L110 sidechain rotating out and the M113 sidechain rotating in to replace it. In association with these changes, the two buried structured waters observed in the bound form also adopt new position within the CDR3 loop, which help to stabilize the new conformation by forming 2 water mediated interactions between the amide backbones of Q109 and G98 with the D36 sidechain, respectively (**figure S6**). In both simulations of the bound and unbound forms of VH-DiF^CAP^-01 (**figure S7**), the two water positions remained predominately solvated indicating that water molecules found here exchanged slowly with the external solvent and formed enthalpically favorable interactions.

### VH-DIF^CAP^-01 Inhibits eiF4F Mediated Cap Dependent Translation by Disrupting the Interplay Between eIF4E and Capped-mRNA

As predicted from in vitro studies, FLAG-tagged VH-DiF^CAP^-01 immuno-precipitated cellular eIF4E more efficiently than VH-DIF^CAP^-02 (**Figure 5A**), a peptide aptamer that demonstrate little binding with eIF4E in vitro (**table 2**). Consistent with the ability of VH-DIF^CAP^-01 to interact with eIF4E at the cap-binding site, eIF4E also co-immunoprecipitated with endogenous eiF4G and 4EBP1 (**Figure 5A**) This result correlated with the reduction of eIF4F complex formation on m^7^GTP beads when VH-DIF^CAP^-01 is over-expressed (**Figure 5B**). Concomitantly with VH-DIF^CAP^-01 expression, a reduction in the levels of eIF4E phosphorylation at S209 was observed (**Figure 5C**). This decrease in eIF4E phosphorylation implies that the VH-DIF^CAP^-01 interaction with eIF4E interfered with the eIF4G mediated recruitment of the MNK1/2 kinase to the eIF4F complex and in turn targeting of the S209 residue.^31^ Mutation of the methionine residue (M104) to alanine, critical for the interaction of the cyclic peptide motif with eIF4E, abrogated the ability of VH-DIF^CAP^-01 to immuno-precipitate eIF4E and confirmed its specific effects on eIF4E phosphorylation. These results infer that either displacement of mRNA from the cap-binding site or steric occlusion caused by VH-DIF^CAP^-01 binding prevents MNK1/2 mediated phosphorylation of eIF4E. The effects of VH-DIF^CAP^-01 upon mRNA translation were assessed using a bi-cistronic luciferase reporter^32^, which demonstrated that the peptide aptamer specifically inhibited cap-dependent translation versus cap-independent translation (**Figure 5D**). Additionally, cellular expression of VH-DIF^CAP^-01 also down-regulated Cyclin D1 protein levels (**Figure 5C**). A protein that is considered to a hallmark to eIF4F signalling inhibition in cells.^33,34^. In contrast, the VH-DIF^CAP^-01 mutant (M104A, “VH-DIF^CAP^-01 MA”) constructs exhibited negligible activity in the bicistronic assay or on cyclin D1 protein expression (**Figure 5C and 5D**). Purified VH-DIF^CAP^-01 was also able to efficiently interact with both phosphorylated and unphosphorylated forms of eIF4E (**Figure 5E**) in pull downs from cell lysate. Examination of the crystal structure demonstrates that phosphorylation of 209 would not impede the eIF4E: VH-DIF^CAP-01^ interaction (**Figure 4I**).

**Figure 5:**
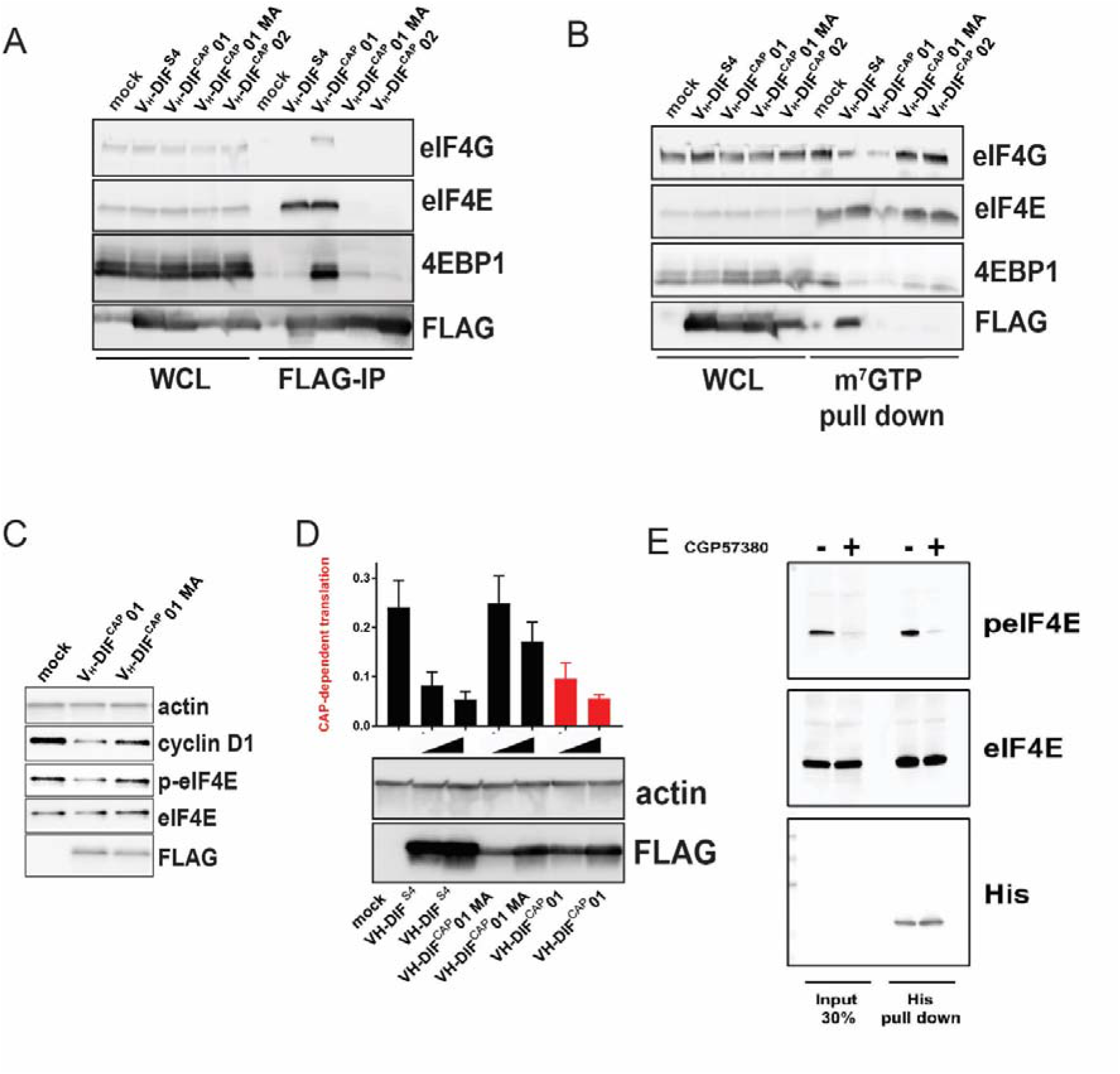
**A)** Anti-FLAG IP pull down of HEK293 cells transfected with either VH-DIF^CAP^-01, VH-DIF^CAP^-02, VH-DIF^CAP^-Cntrl and VH-S4, a VH-domain that interacts with eIF4E at the eIF4G binding interface. IP experiments were performed 24 hours post transfection. Whole cell lysate (WCL) was also blotted for the corresponding proteins and is shown on the left of the blot. **B)** m^7^GTP pulldown of eIF4E containing complexes from HEK293 transfected with VH-DIF^CAP^01, VH-DIF^CAP^02, VH-DIF^CAP^01 MA (M104A) and VH-S4. Whole cell lysate (WCL) was also blotted for the corresponding proteins and is shown on the left of the blot. In the blot below an equivalent pull-down was performed but with the HEK293 cells transfected with increasing amounts of expression vector. **C)** HEK293 cells were transfected with either empty vector, VH-DIF^CAP^01 or VH-DIF^CAP^01 MA (M104A), and eIF4E phosphorylation and cyclin D1 expressions levels assessed via western blot. Actin was used as a loading control, whilst anti-FLAG was used to assess expression of the transfected proteins. Protein levels were assessed 48 hours post transfection. **D)** A bicistronic luciferase reporter, which measures the relative amount of cap-dependent translation (Renilla) to cap-independent translation (Firefly), was co-transfected with either empty vector (MOCK) or increasing amount of VH-DIF^CAP^-01, VH-DIF^CAP^-01 MA, VH-S4 plasmid vector into HEK293 cells (see material and method). Renilla and Firefly luciferase activity was measured 48⍰h post transfection and plotted as a ratio-metric value. **E)** Anti-His IP pulldown of purified VH-DIF^CAP^-01 exogenously added to HEK293 cell lysates either treated with CGP57380 or vehicle control. Input lysate is shown on left hand side of the western blot.

### Development of a Novel Live Cell Protein-Protein Interaction (PPI) Assay to Measure Antagonism of the eIF4E Cap-Binding Site

Currently, there is no live cell-based assay that can evaluate engagement of the cap-binding site by small molecules or other modalities. A site that has been the target of multiple studies to develop cell permeable small molecules for therapeutic development. Fortunately, there is a plethora of suitable technologies that can be used to develop an appropriate assay e.g., split luciferase^35^, BRET^36^ and FRET^37^ (**<u>b</u>**io and **<u>f</u>**luorescence **<u>r</u>**esonance **<u>e</u>**xcitation **<u>t</u>**ransfer), and cellular localization technologies.^38^ We therefore sought to exploit the VH-DIF^CAP^-01 peptide aptamer in combination with the NanoLuc-based protein complementation system (NanoBit, PCA, Promega)^36^ to develop a PPI assay that can assess antagonism of the m^7^GTP cap-binding site in eIF4E in cells. The NanoLuc complementation protein system consists of two components termed LgBiT (18-kDa protein fragment) and SmBiT (11-amino-acid peptide fragment), which have been optimised for minimal self-association and stability. When LgBiT and SmBiT are optimally fused to two interacting proteins, they will be brought into proximity to each other by the resulting interaction, resulting in the formation of the active luciferase.

The LgBIT-eIF4E and SmBIT-VH-DIF^CAP^-01 were identified as the transfection pair that reconstituted the highest luciferase signal without exhibiting background activity in the negative controls, thus confirming that the NanoBit reporter fragments were not spontaneously assembling under the experimental conditions (**Figure 6A and 6B**). To further confirm the specificity of this assay for binding at the cap site, the NanoBit assay was re-performed with both the VH-DIF^CAP^-01^M104A^ (termed VH-DIF^CAP^-01 “MA”) and VH-DIF^CAP^-01^E103A^ (termed VH-DIF^CAP^-01 “EA) binding controls fused to smBIT, which resulted in the abrogation of the luciferase signal above background (**Figure 6B**). Additionally, the ability of the assay to measure and differentiate between interactors that bound at the either the cap-binding or eIF4G binding sites of eIF4E was demonstrated by co-transfection of the NanoBit assay (termed NanoBIT eIF4E^CAP^) with either untagged VH-M4, a VH-domain designed to interact with eIF4E at the eIF4G binding site, or VH-DIF^CAP^-01 (**Figure 6C**).

**Figure 6:**
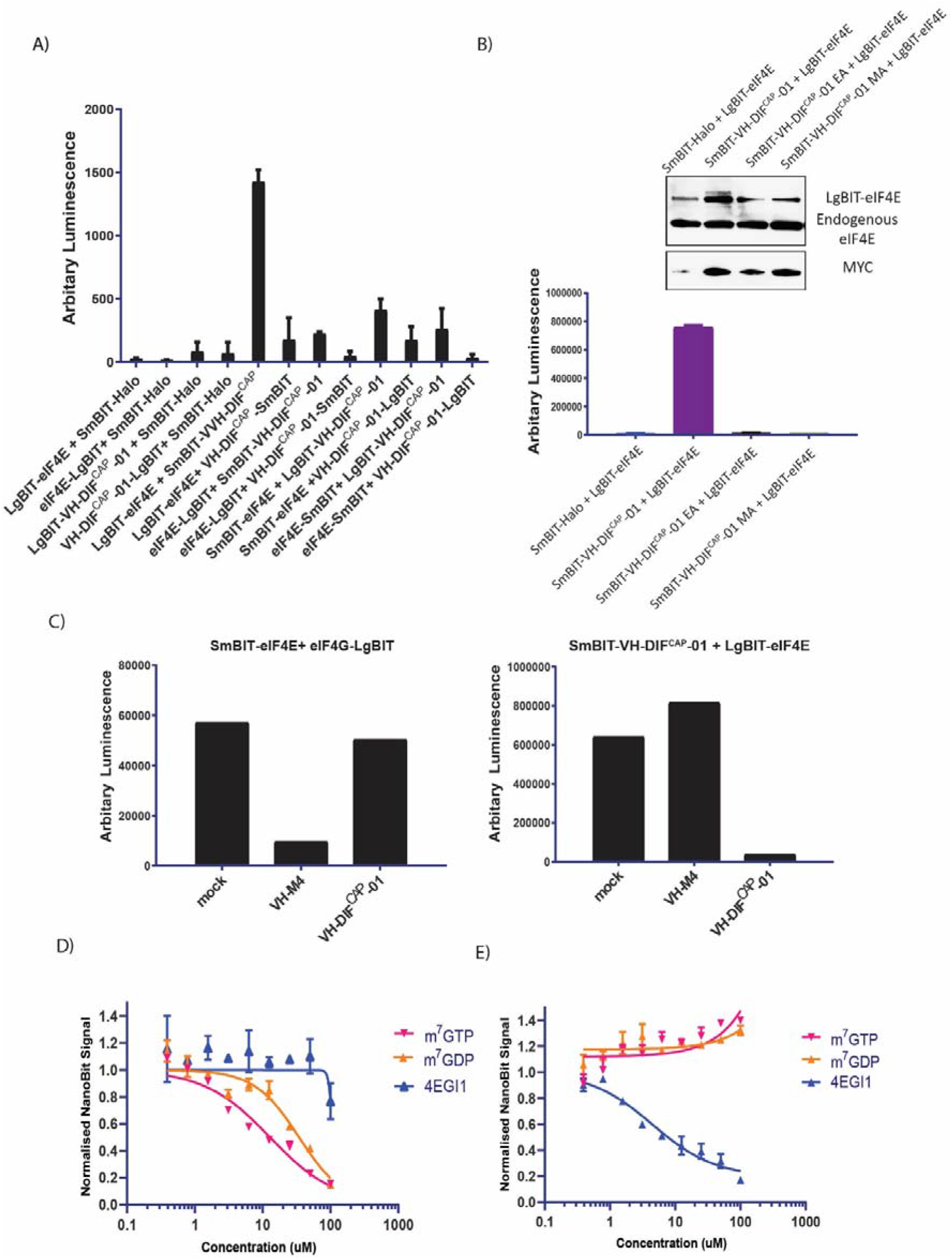
**A)** Inset shows how the interaction of proteins A and B fused to SmBiT and LgBiT (Promega, USA) enables reconstitution of the active NanoBit (Promega, USA) luciferase. Graph shows the reconstituted luminescence activity of the various combinations of either eIF4E or VH-DIF^CAP^-01 fused at either the N or C terminal of SmBiT and LgBiT, respectively, co-transfected into HEK293 cells. Individual N- and C-terminal LgBiT-linked eIF4E and VH-DIF^CAP^-01 constructs co-transfected with SmBiT-HALO served as negative controls. **B)** To validate the specificity of the SmBIT-VH-DIF^CAP^-01 and LgBiT-eIF4E interaction pair, two VH-DIF^CAP^-01 point mutant controls were generated (E103A (EA) and M104A (MA), respectively) and co-transfected into HEK293 cells with LgBiT eIF4E, which resulted in loss of bioluminescence. **Inset**: Cell samples replicating the NanoBiT experimental conditions were assessed for their relative levels of LgBIT fused eIF4E to endogenous eIF4E, and expression levels of the various SmBIT-VH-DIF^CAP^-01 constructs. **C)** (**Right hand graph**) The ability of the SmBIT-VH-DIF^CAP^-01: LgBiT-eIF4E (termed NanoBIT^CAP^) interaction pair to discriminate between different classes of eIF4E binders was tested by co-expressing it with either VH-S4 (a VH domain that interacts specifically with the eIF4G interaction site) or VH-DIF^CAP^-01 not fused to SmBIT, where only VH-DIF^CAP^-01 caused a decrease in luminescence. (**Left hand graph**) The specificity of VH-DIF^CAP^-01 was further investigated by co-expressing either VH-DIF^CAP^-01 or VH-S4 with the NanoBit eIF4E:eIF4G6^04–646^ system, which measures binding at the eI4G interface and demonstrated that VH-DIF^CAP^-01 only interacts with the cap-binding interface. **D)** HEK293 cells were transfected with the NanoBIT^CAP^ system and permeabilized with a sub-CMC (critical micelle concentration) concentration of digitionin. Cells were then treated with different titrations of small molecules that either specifically targeted the cap (m GTP, m GDP) or eIF4G (4EGI) binding interfaces of eIF4E. **E)** HEK293 cells were transfected with the NanoBit eIF4E:eIF4G604–646 system and again permeabilized with a sub-CMC (critical micelle concentration) concentration of digitionin. Cells were then treated with titration of the following compounds (m^7^GTP, m^7^GDP and 4EGI) to assess the specificity of the NanoBIT^CAP^ system.

To validate that the NanoBIT eIF4E^CAP^ system can measure small molecule mediated inhibition of the eIF4E cap-binding site, the system was also used to screen two known cap-analogue antagonists (m^7^GTP and m^7^GDP) and an established cell permeable inhibitor of the eIF4E:4G interface (4EGI1) as a negative control. Unfortunately, both cap-analogue molecules are cell impermeable. Therefore, to circumvent this issue, a sub-CMC (critical micelle concentration) treatment of digitonin was used to permeabilize and enable cellular entry of the cap-analogues into HEK293 cells transfected with the NanoBIT^CAP^ system. In the permeabilized cells, both m^7^GTP and m^7^GDP disrupted the interaction of LgBIT-eIF4E with SmBIT-VH-DIF^CAP^-01 with IC_50_ s of 12.8 μM and 34.5 μM respectively, whilst 4EGI1 had a negligible effect on the NanoBIT signal, demonstrating that the assay system can measure cap-binding site antagonists (**Figure 6D**). To highlight the specificity of the NanoBIT^CAP^ system further, it was also shown that only 4EIG1 and neither of the two cap-analogues ^7,8^ were able to inhibit the luciferase signal in an alternative NanoBIT system (eIF4E:eIF4G604–646) that measures disruption of the eIF4E:4G interface (**Figure 6E**). Both sets of described experiments were then repeated in non-permeabilized cell, where as expected the impermeable cap-analogues elicited no effects, and the cell permeable only 4EGI1 inhibited the NanoBIT signal in the NanoBit eIF4E:eIF4G^604–646^ system.

## Conclusion

A novel cyclic peptide (EE-02) was identified that interacts with eIF4E, a target of high therapeutic relevance in oncology, through its mRNA binding site via an unreported binding pose. To enable further study of this antagonist upon the cellular eIF4E:Cap interaction and to overcome the inherent liabilities of the cyclic peptide (intracellular reduction of the disulphide bond, proteolytic instability, and lack of cell permeability) preventing its use in cells, the critical residues of the EE-02 binding epitope were evolved on to the CDR3 loop of a suitable monomeric VH-domain through a process termed Peptide Epitope Linker Evolution (PELE), resulting in the development of the cap-site interacting peptide aptamer VH-DIF^CAP^-01. The PELE system entails the insertion of the defined EE-02 binding ‘epitope’ into a the CDR3 loop region of the VH domain and the residues either side of the motif (the ‘linkers’) in the CDR3 loop evolved to recapitulate the conformation required for binding eIF4E. We additionally demonstrated that the VH-DIF^CAP^-01:eIF4E complex structurally recapitulated the EE-02 interaction, could be intracellularly expressed and that it could also modulate known biological activities of eIF4E.

An interesting feature of the presentation of the EE-02 binding motif in VH-DiF^CAP^-01 is the role played by two buried structured waters that facilitate the transition of the VH domain from its unbound form to the bound form with eIF4E. These waters in combination with the linkers isolated in the yeast selection allow the CDR3 loop to adopt the β-hairpin structure that the “EMGFF” epitope requires to bind to eIF4E and to accommodate the D38:R51 salt-bridge that the CDR3 loop sit upon. The CDR3 loop of VH-DiF^CAP^-01 folds back onto the former light-chain interaction surface, where the D38:R51 salt-bridge is located, in a manner highly like that seen in nanobodies (**figure S8**). Also like many other VH and VHH structures and their interaction with their binding partner, the interaction with eIF4E is principally mediated through the CDR3 loop. However, this type of interaction does differ significantly from the reported VH domain interaction with VEGF, where both the CDR3 and the the former light-chain interaction surface are involved in macromolecular recognition (**figure S9**). The identified cap-binding site interaction pose can also be used as a template for the development of new therapeutic modalities. Furthermore, to support the discovery of drug-like cap-analogue molecules we have also used the VH-DiF^CAP-01^ peptide aptamer to develop a split-luciferase assay that can assess cellular target engagement of the cap binding site in live-cells. In addition, VH-DiF^CAP^-01 can also be used to further study the biological role of the eIF4F complex, especially in combination with the reported VH-S4 domain that disrupts eIF4F complex formation by binding eIF4E at its 4G interaction site. A suite of tools enabling discrimination of the effects of inhibition at precise sites on eIF4E in cellular studies. This contrasts with tools such as RNAi where mRNA levels are reduced and do not allow the study of specific functional domains within proteins, and in turn any insights may be therapeutically limited.

We also envision that the VH-DIF^CAP^-01 molecule and other such molecules, when combined with new and novel alternate delivery methods e.g., RNA, toxin, and cell-penetrating peptide mediated delivery methods, could also be used to target cap-dependent translation and constitute a potential therapeutic. These results demonstrate that the PELE process can enable the development of mini-proteins that interact at desired target interaction sites, that can model and assess the potential effects of drug inhibition and allow the construction of critical target engagement assays that can accelerate the identification of lead compounds for drug development.

## Notes

### Competing Interest Statement

I.A. is a shareholder of DotBio Pte. Ltd.
I.A. is an employee of DotBio Pte. Ltd.

### Summary of Updates

Minor edit: 1) the introduction of specific or hypervariable loop sequences into scaffolds that are not optimized for stability can lead to loss of peptide conformation to 2) specific or hypervariable loop sequences are inserted into scaffolds that are not optimized to stabilise their bound conformations

