## Supplemental Table and Methods for "Development of a Novel Peptide Aptamer that Interacts with the eIF4E Capped-mRNA Binding Site using Peptide Epitope Linker Evolution (PELE)"

**Supplementary Information**

| **PDB ID:** | 7EZW | 7F07 |
| --- | --- | --- |
| **Resolution (Å)** | 48.148- 2.35, (2.47-2.35) | 40.996-2.25, (2.37-2.25) |
| **Space Group** | P6_3_ | P2_1_2_1_2_1_ |
| **Unit Cell Dimensions (Å)** | a = 81.14, b = 81.4, c = 66.11, α = γ = 90°, β = 120° | a = 38.52, b = 81.82, c = 122.99, α = β = γ = 90° |
| **Temp (K)** | 100 | 100 |
| **Redundancy** | 11.3, (11.2) | 13.4, (14.0) |
| **Unique Collected Reflections** | 10511, (1027) | 19,226, (2734) |
| **Completeness (%)** | 99.9, (99.2) | 100, (100) |
| **R Merge (%)** | 0.114, (0.711) | 0.120, (0.755) |
| **I/sigma** | 13.8, (2.6) | 4.7, (1.0) |
| **R factor (%)** | 20.53 | 22.19 |
| **R free (%)** | 24.76 | 27.18 |
| **RMS Bonds (Å)** | 0.0023 | 0.0028 |
| **RMS Angles (°)** | 1.143 | 1.216 |
| **Wilson B-factor (Å^2^)** | 30.5 | 37.6 |
| **Average Refined B Factors** |  |  |
| **Chain A** | 35.29, (eIF4E) | 48.34, (eIF4E) |
| **Chain B** | 41.85, (EE-02) | 56.6, (VH-DiF^CAP^-01) |
| **Waters** | 28.67 | 46.46, |
| **Number of Water Molecules** | 59 | 48 |
| **Ramachandran Data (Rampage). Number of Residues in (%):** |  |  |
| **Favoured Region** | 96.7 | 97.8 |
| **Allowed Region** | 3.3 | 2.2 |
| **Outlier Region** | 0 |  |

**Table S1:** Crystallographic data collection and refinement statistics. Highest resolution bin data stated in parentheses.

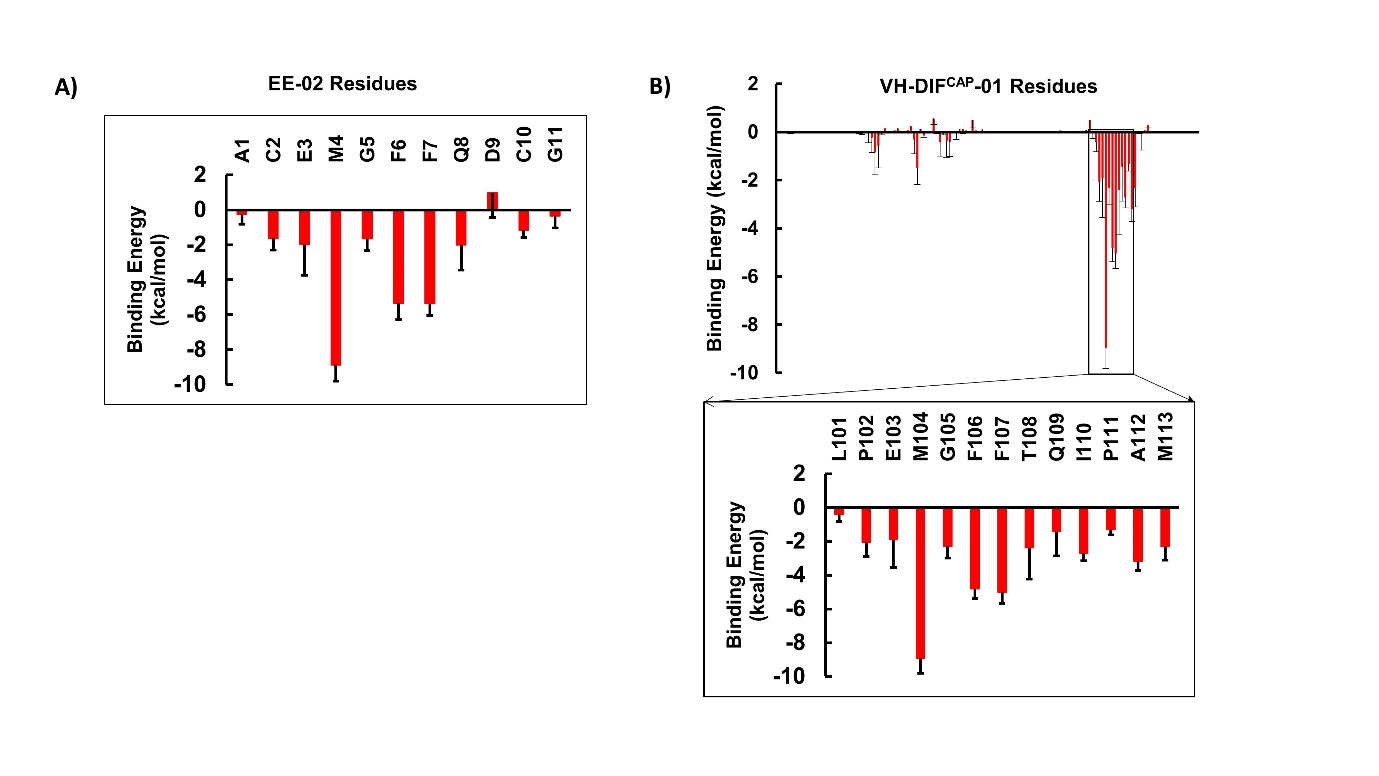

**Figure S1:** Binding energy decomposition analysis from A) MD simulations of the eIF4E:EE-02 complex structure which demonstrates that M4, F6 and F7 contribute a significant proportion of the binding energy of the complex, and B) MD simulations of the eIF4E:VH-DIF^CAP-01^ complex showing that the EE-02 motif also underpins the energetics of the VH domain’s interaction with eIF4E. The enlarged portion of the graph details the precise contributions being made by each residue of the interaction motif to the binding energy. Again, residues F106, F107 and M104 (equivalent to M4, F6 and F7 in EE-02) contribute significantly to the energetics of the complex.

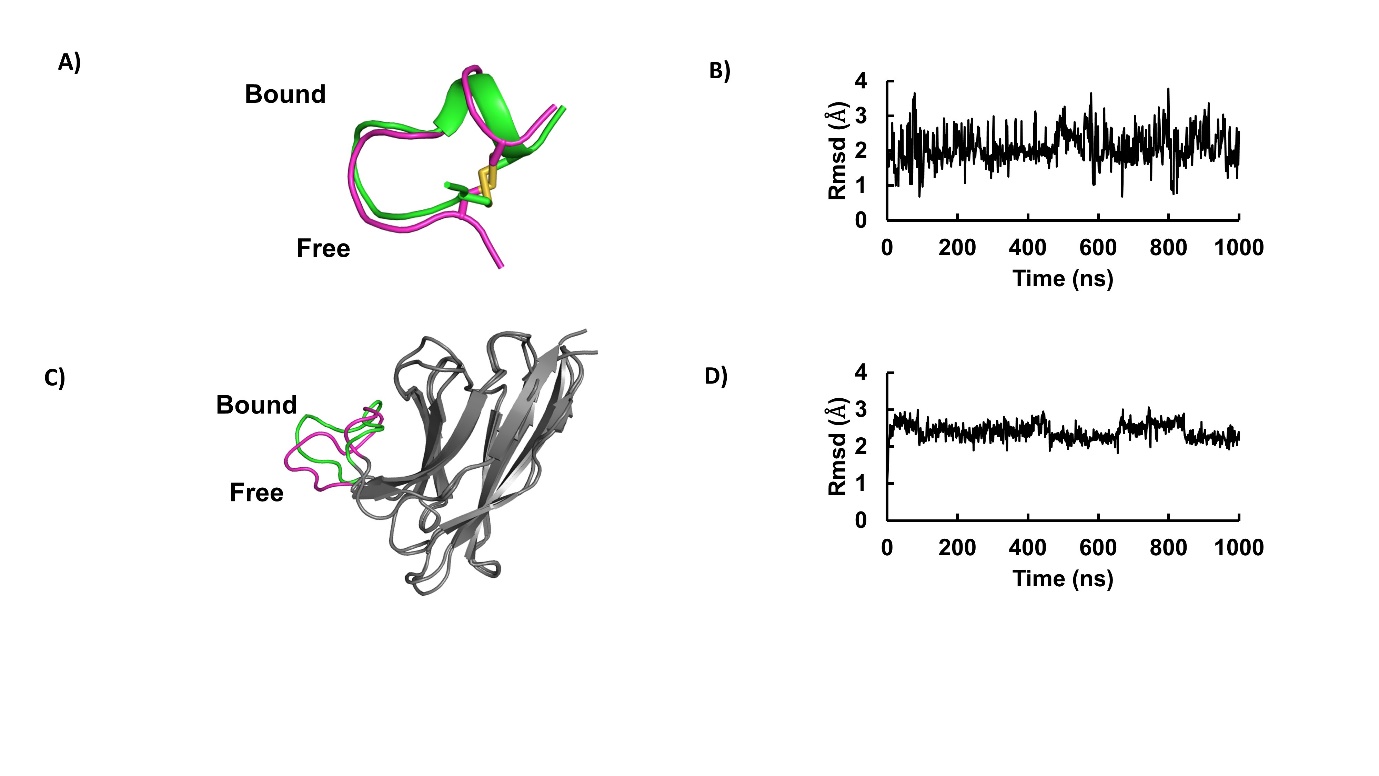

**Figure S2:** Structural deviation between bound and free states of VH-DIF^CAP^-01 and EE-2. A) The averaged free state EE-02 structure, derived from MD simulations of the cyclic peptide alone, overlaid with the bound structure of EE-02 from the eIF4E:EE-02 crystal structure. B) RMSD plot of the MD simulation frames of unbound EE-02 against the bound crystal form. C) The averaged free state VH-DIF^CAP^-01 structure, derived from MD simulations of the VH domain alone, overlaid with the eIF4E bound structure of VH-DIF^CAP^-01 from the crystal structure. The RMSD values sample a broad range inferring that the EE-02 peptide adopts a fold similar to the bound form, but is relatively flexible. D) RMSD plot of the MD simulation frames of unbound VH-DIF^CAP^-01 against the bound VH domain crystal form.

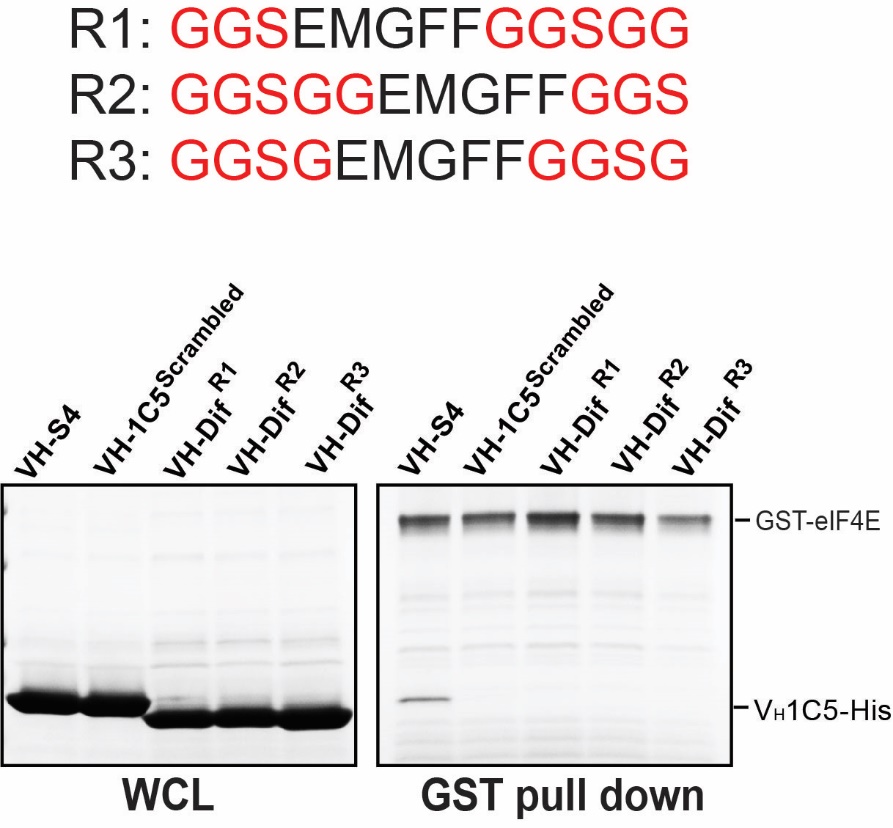

**Figure S3:** His-Tagged VH constructs (R1, R2, R3) with the cap-site binding peptide motif rationally grafted on at alternate positions (see insert) in the CDR3 loop were screened in a pull-down assay against glutathione beads with bound GST–tagged eIF4E. The left-hand panel shows the protein input into the assay, whilst the right-hand panel shows the results of the pull-down after stringent washing. VH-1C5, a VH-domain that has been shown to interact at the eIF4E:4G interface was used as a positive control, whilst VH-1C5^Scrambled^, where the corresponding CDR3 loop has been scrambled, was used as a negative control.

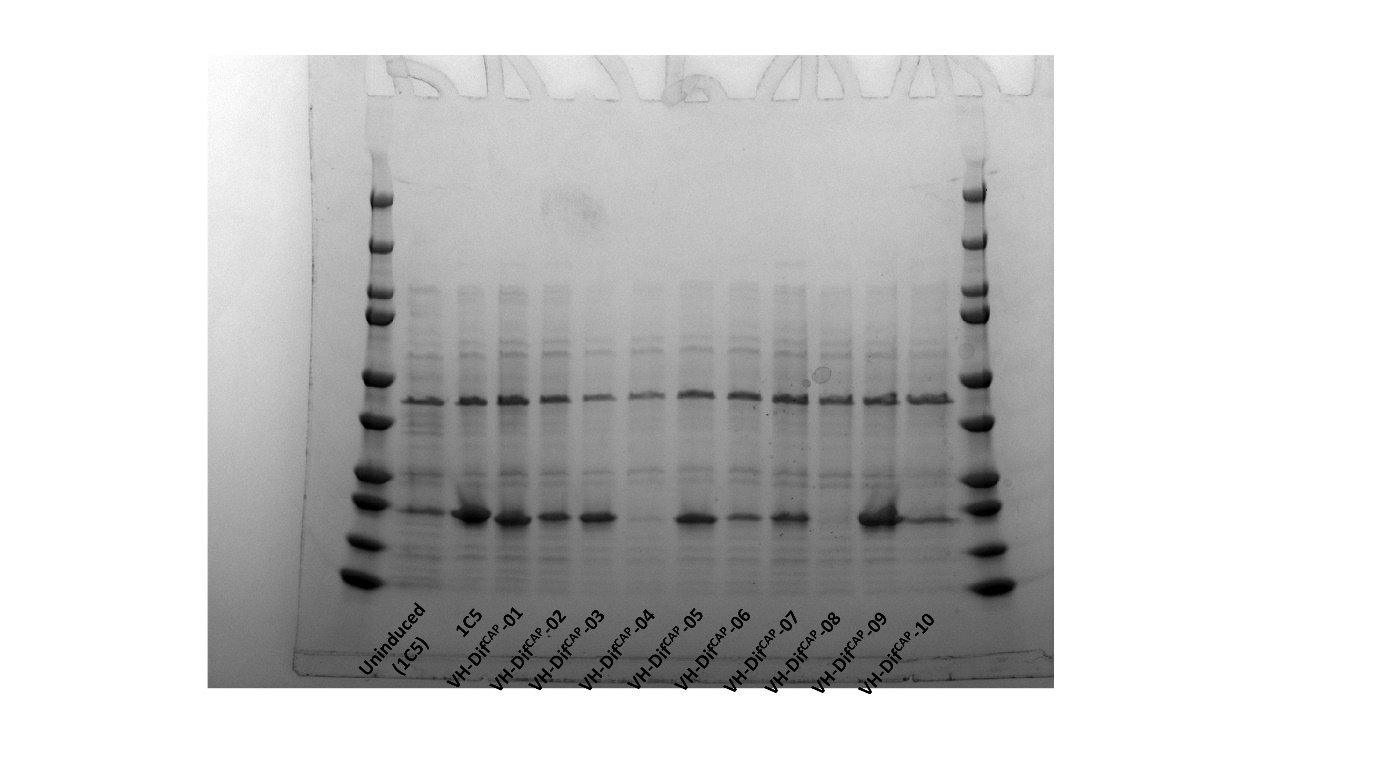

**Figure S4:** The VH-DiF^CAP^ peptide aptamers identified by the yeast-based peptide epitope linker evolution experiments were tested for soluble expression in small scale bacteria cultures. His-tagged proteins were purified using Ni^2+^ chelated IMAC spin columns and analyzed using coommasiie stained SDS PAGE gels to detect soluble protein. VH-Dif^CAP-01^, VH-Dif^CAP-05^ and VH-Dif^CAP-09^ were selected for scaling up and further interaction analysis. The previously characterized VH domain (IC5), which interacts with eIF4E at the 4G binding site was used as a positive control.

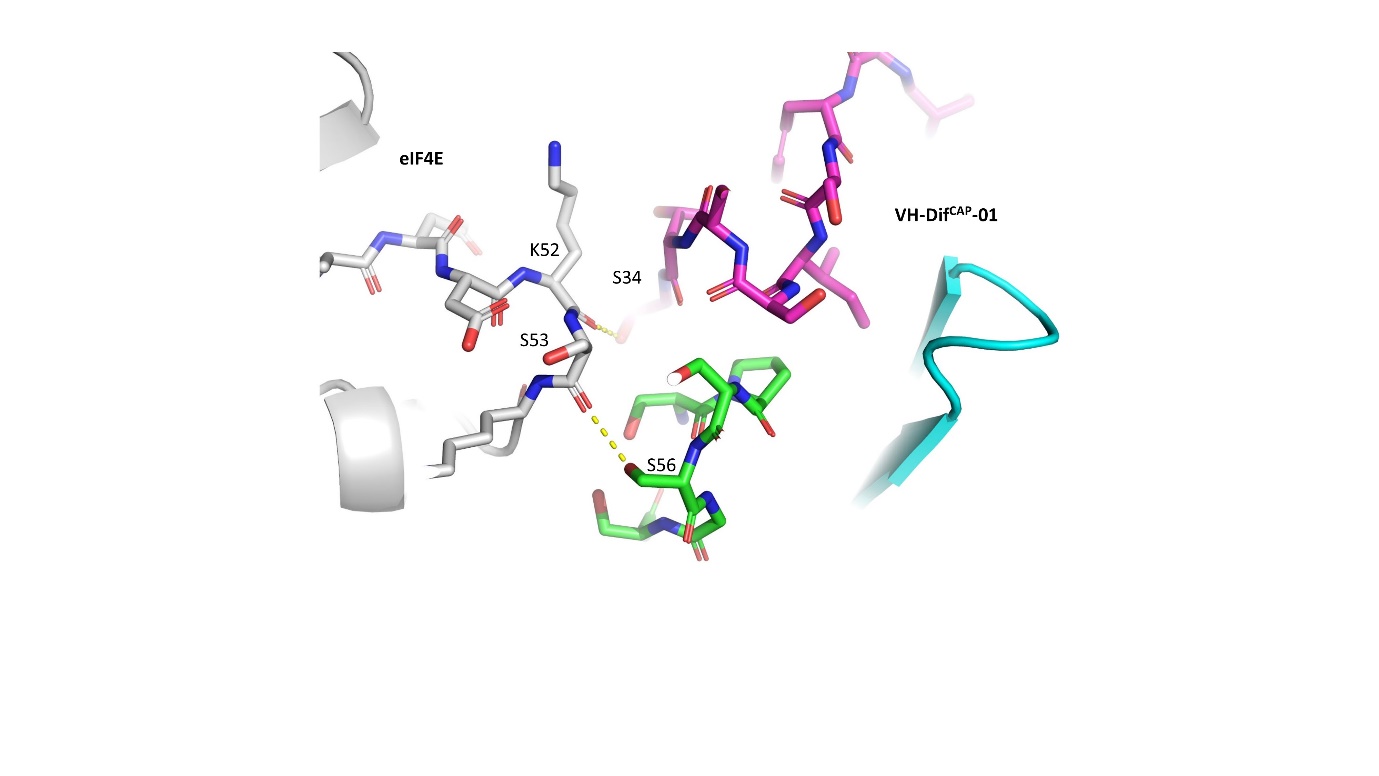

**Figure S5:** VH-DiF^CAP^-01 also interacts weakly at a second site with eIF4E (white) that is constituted from the CDR1 (magenta) and CDR2 (green) loops of the VH domain, where the residues S34 and S56 both form hydrogen bonds via their sidechains with the carbonyls of K52 and S53, respectively.

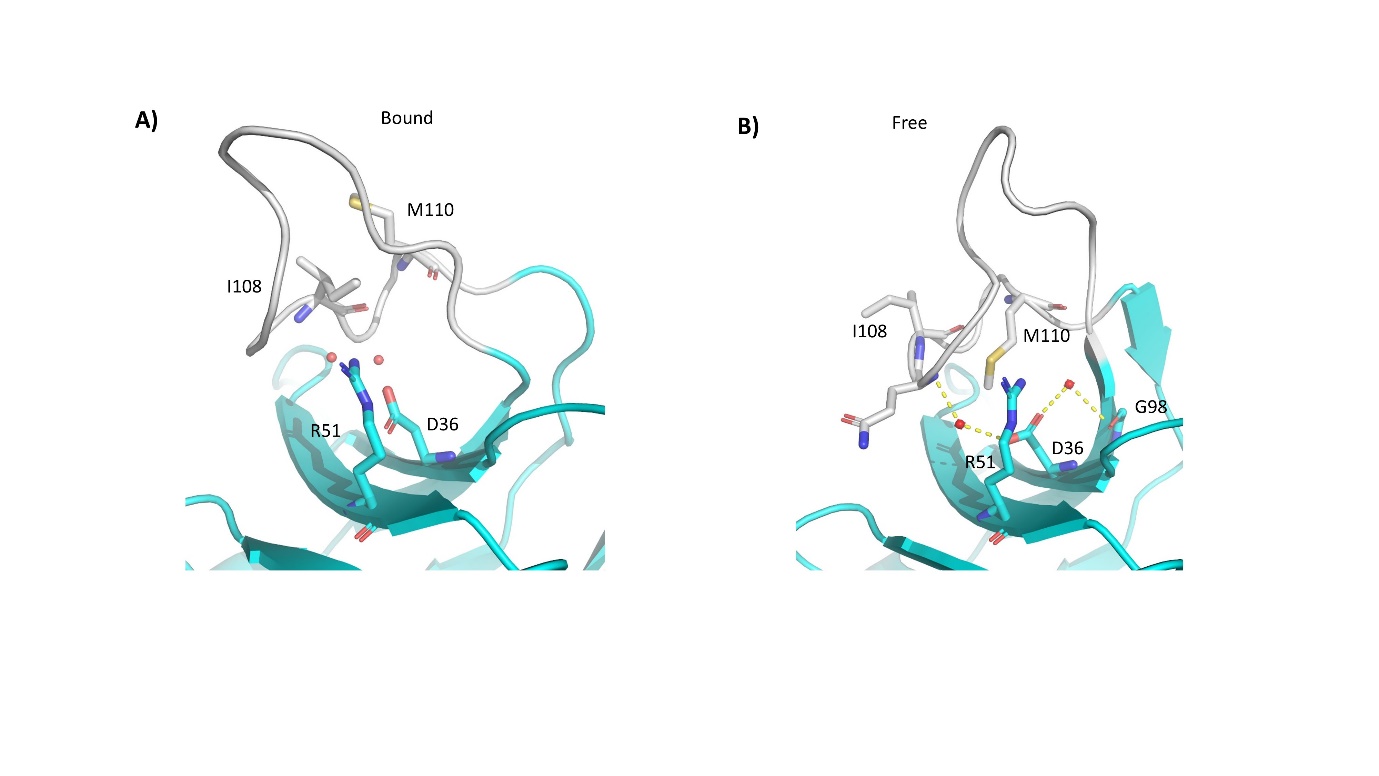

**Figure S6: A)** Crystal structure of VH-DiF^CAP^-01 bound to eIF4E. Buried structured waters are depicted with red spheres. The CDR3 loop bearing the ‘EMGFF’ cap-binding site interaction motif is highlighted in white. B) Averaged structure of unbound VH-DiF^CAP^-01 derived from MD simulations (see materials and methods). The CDR3 loop undergoes a structural relaxation, whereby the β-hairpin structure associated with the ‘EMGFF’ motif is lost and instead there is a general movement of the CDR3 loop away from the body of the VH domain scaffold. Interestingly, this movement is underpinned by significant structural changes in the hydrophobic core of the CDR3 loop structure with the L110 sidechain rotating out and the M113 sidechain rotating in to replace it. In association with these structural re-arrangements, the two buried structured water observed in the bound form (red spheres) also adopt new position within the CDR 3 loop, which help to stabilize the new conformation by forming 2 water mediated interactions (dashed yellow lines) between the amide backbones of Q109 and G98 with the D36 sidechain, respectively

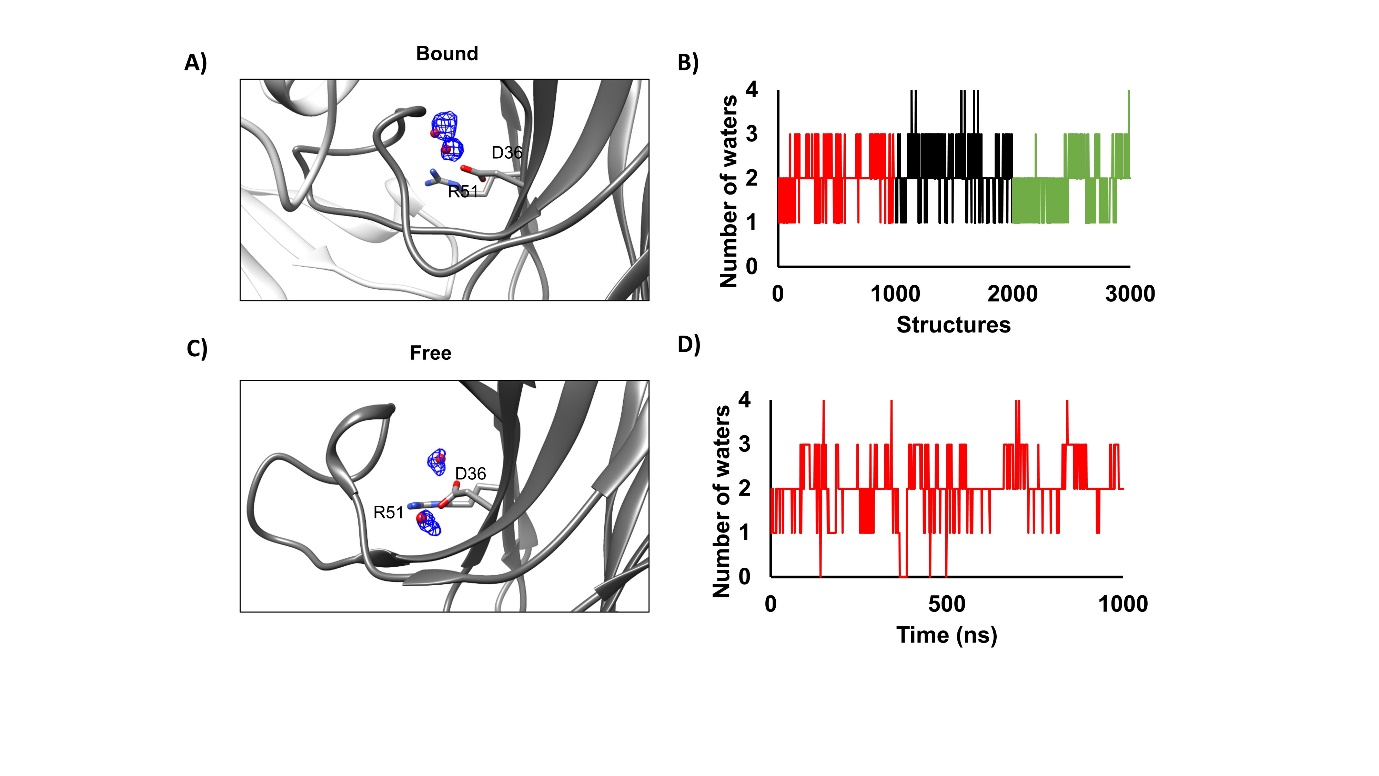

**Figure S7:** Solvation properties of free and bound VH-DIF^CAP^-01: A) Solvation map depicting the averaged positions of the two buried waters, derived from the MD simulations of VH-DIF^CAP^-01 in complex eIF4E, that stabilize the CDR3 loop conformation. B) Graph depicting the number of water molecules observed at and within the vicinity of the 2 average water molecule position shown in A). C) Solvation map depicting the averaged positions of the two buried waters, derived from the MD simulations of unbound VH-DIF^CAP^-01, that change their position significantly in relationship to the bound form. D) Graph depicting the number of water molecules observed at and within the vicinity of the 2 average water molecule position shown in C).

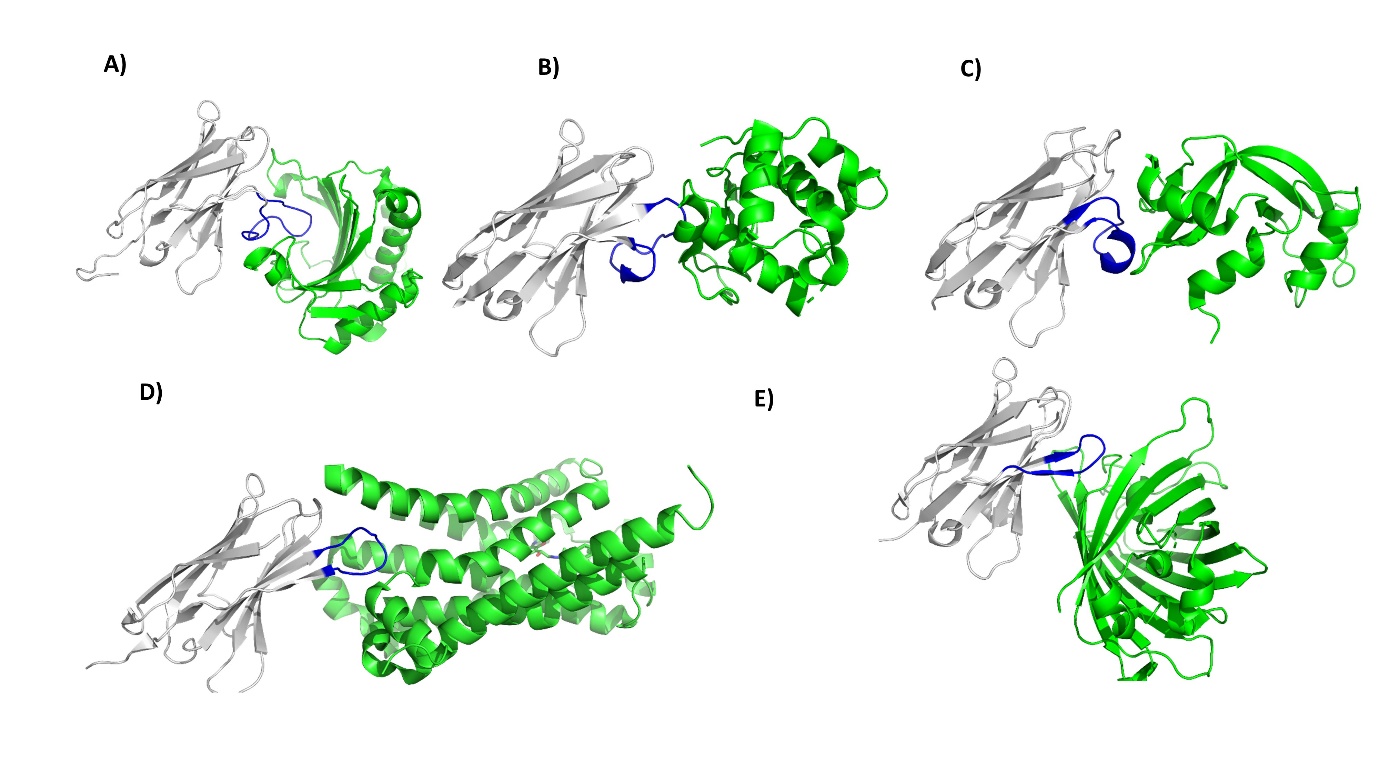

**Figure S8:** A) The CDR3 loop (blue) of VH-Dif^CAP^-01 (white) folds back onto the former light-chain interaction surface of the VH domain. B) NanoBodies (VHH domain derived from a camelid antibody) in complex with lysozyme (PDB ID: 1Z4H) and C) RNase A (PDB ID: 2P4A) showing the interacting CDR3 loops (blue) folding back onto the main body of the VHH domain (white). Nanobodies in complex with D) β2 adrenoceptor (adrenoceptor-PDB ID: 3P0G) and E) GFP (PDB ID: 3K1K), where the CDR3 interacting loops form no packing interactions with the VHH domains themselves.

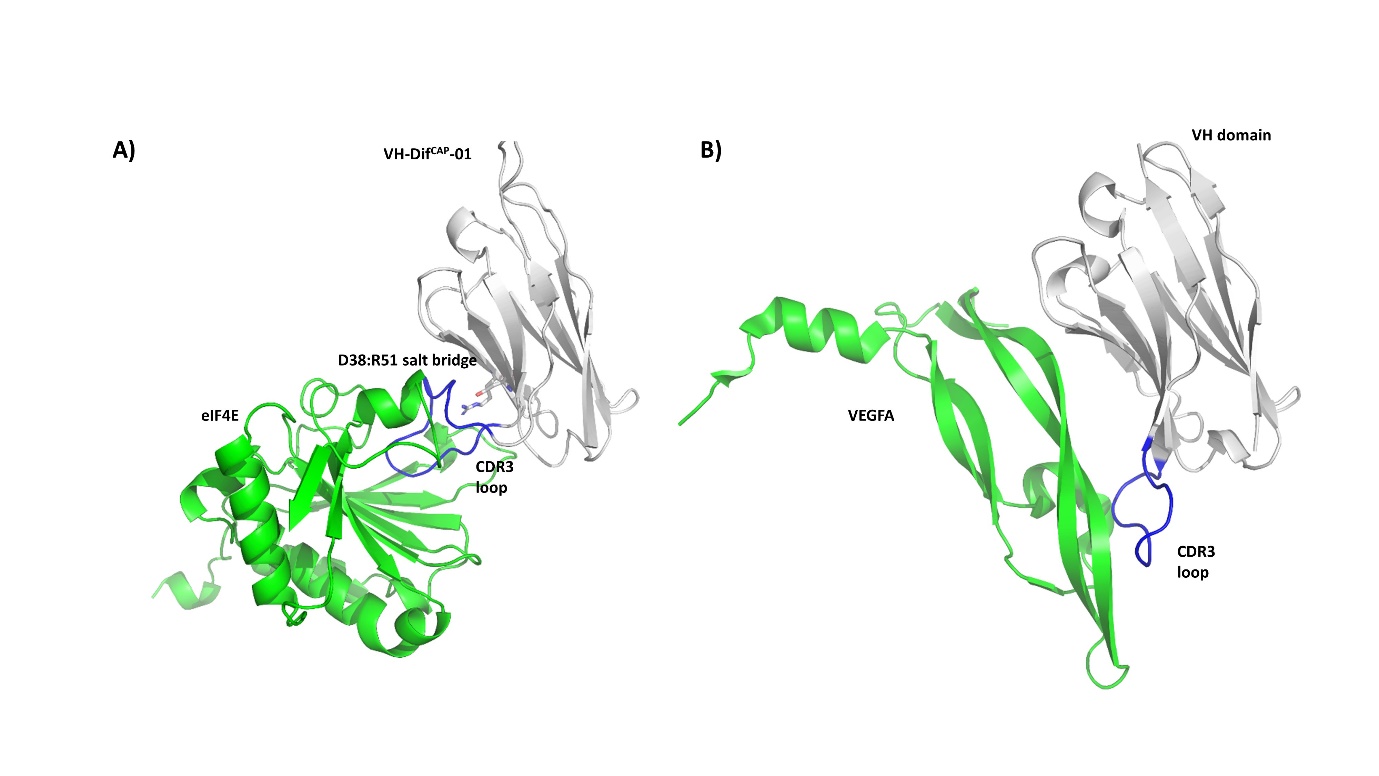

**Figure S9:** A) The CDR3 loop (blue) of VH-Dif^CAP^-01 (white) folds back onto the former light-chain interaction surface, where the D38:R51 salt-bridge is located. The interaction of the CDR3 loop (blue) with the salt-bridge stabilizes its conformation enabling it to engage the cap-binding site on eIF4E (green). B) However, this type of interaction does differ significantly from the reported VH domain interaction with VEGFA (PDB ID: 3P9W), where both the CDR3 (blue) and the the former light-chain interaction surface are involved in macromolecular recognition. It also must be noted the CDR3 loop does not fold back on to the VH domain.

**Methods and Material**

**Peptide Synthesis**

**Peptides** were ordered from and synthesized by Mimotopes, Clayton, Australia. Peptides were purified using HPLC to >90% purity. All peptides were amidated at their C-terminus and acetylated at their N-terminus. Peptides were purified using HPLC to >90% purity.

**eIF4E expression and purification for crystallisation and biophysical assays.**

Full-length human eIF4E was expressed and purified as described previously.^1^

**eIF4E expression and purification for Sortase Labelling**

eIF4E was cloned into the GST fusion expression vector pGEX-6P1 (GE Lifesciences). BL21 DE3 competent bacteria were then transformed with the GST-tagged fusion constructs. A single colony was picked and grown in LB medium at 37 °C to an OD600 of ~ 0.6 and induction was carried out overnight with 0.3 mM IPTG at 16 °C. Cells were harvested by centrifugation, and the cell pellets were resuspended in PBS (Phosphate Buffered Saline, 2.7 mM KCl and 137 mM NaCl, pH 7.4) and then sonicated. The sonicated sample was centrifuged for 60 min at 17,000*g* at 4 °C. The supernatant was applied to a 5 ml FF GST column (Amersham) pre-equilibrated in PBS buffer with 1 mM DTT. The column was then further washed by 6 volumes of PBS. Proteins were then purified from the column by either **1)** cleavage with thrombin (Sigma-Aldrich) protease or **2)** elution with glutathione.

**1)** Ten units of thrombin (Sigma-Aldrich) protease, in one column volume of PBS with 1 mM DTT buffer, were injected into the column. The cleavage reaction was allowed to proceed overnight at 4 °C. The cleaved protein was then eluted off the column with wash buffer. Protein fractions were analysed with SDS page gel and concentrated using a Centricon (3.5 kDa MWCO) concentrator (Millipore). Protein samples were then dialyzed into a buffer solution containing 20 mM Tris pH 8.0 with 1 mM DTT and loaded onto a mono Q column pre-equilibrated in buffer A (20 mM Tris, pH 8.0, 1 mM DTT). The column was then washed in 6 column volumes of buffer A and bound protein was eluted with a linear gradient of 1 M NaCl over 25 column volumes. Protein fractions were analysed with SDS page gel and concentrated using a Centricon (3.5 kDa MWCO) concentrator (Millipore). The cleaved constructs were then purified to 90% purity. Protein concentration was determined using A280.

**2)** The column was then washed in 6 column volumes of PBS with 1 mM DTT and bound protein was eluted with a linear gradient of 50 mM Tris, pH 8.0, 0.1 M NaCl with 250 mM glutathione over 10 column volumes. Protein fractions were analysed with SDS page gel and concentrated using a Centricon (3.5 kDa MWCO) concentrator (Millipore). Concentrated protein was dialysed into PBS with 1 mM DTT and protein concentration determined using A280.

**Sortase (SrtA^8M^) Expression and Purification**

The protein sequence corresponding to 61-206 of SrtA *(*Staphylococcus aureus) containing the following mutations (P94R, D160N, D165A, K190E, K196T, E105K, E108A and G167E) was ordered as a gene fragment from IDT (Integrated DNA technologies). The sequence was PCR amplified and inserted into a pNIC-CH bacterial expression plasmid via ligation independent cloning in frame with a C-terminal 6xHis tag. The pNIC-CH-(61-206)SrtA^8M^ (termed SrtA^8M^) expression vector was transformed into BL21(DE3) Rosetta competent cells and a single colony was used to inoculate a 20 ml starter culture in TB (terrific broth containing 25ug/ml of chloramphenicol and 20ug/ml *of* kanamycin), which was incubated overnight at 37°C and shaken at 200 rpm*. The* starter culture was used to inoculate 750 ml of TB and was incubated at 37°C until a O.D_600_ reading of 2.0 was attained. Next, the temperature of the culture was lowered to 18°C and protein expression induced with 0.5 mM of IPTG overnight. Cells were harvested by centrifugation, and the cell pellets were resuspended in 20 ml of lysis buffer (100 mM HEPES pH 8.0, 500 mM NaCl, 10 mM Imidazole, 10 % glycerol, 0.5 mM TCEP, 1000u Benzonase (Merck)) and then sonicated. *The* sonicated sample was centrifuged for 30 min at 17,000*g* at 4 °C. Supernatants were then filtered through 1.2 μm syringe filters and were loaded onto a Ni-nitrilotriacetic acid (NTA) column, pre-equilibrated with 20 mM HEPES pH 7.5, 100 mM NaCl, and 0.5 mM TCEP. The column was then washed with 5 column volumes of the same buffer containing 10 mM Immidazole. Hexahistidine tagged SrtA^8M^ was then eluted with a 1 M imidazole linear gradient. The protein was further purified by size exclusion chromatography (HiLoad 16/60 Superdex 75 prep grade, Cytiva Lifescience) using a 20 mM HEPES pH 7.5, 300 mM NaCl, 10% (v/v) glycerol, 0.5 mM TCEP buffer. Protein concentration was determined using A280 with an extinction coefficient determined from the primary sequence of the construct determined by ProtPARAM.

**N-Terminal biotin labelling of eIF4E mediated by SrtA^8M^**

Sortase-mediated ligation was used to specifically label eIF4E at the N-terminal with biotin. Cleavage of the GST-fused eIF4E with thrombin leaves a single glycine at the N-terminus. The ligation was carried out with thrombin cleaved eIF4E at 50 µM, SrtA^8M^ at 1 µM, and biotin-KGGGLPET-GG-OHse(Ac)-amide peptide at 200 µM in 200 µL of ligation buffer (50 mM Tris pH 8.0, 150 mM NaCl, 1 mM TCEP). SortaseA61-206/8M contains mutations that increase ligation efficiency and make it calcium-independent^2^. The ligation was incubated at room temperature for 4 hours. SrtA^8M^, which contains a C-terminal 6×His-tag, was removed with Dynabeads His-Tag (cat# 10104D, Thermo Fisher). The biotinylated protein was then dialyzed at 4 °C using slide-A-Lyzer cassette (10k MWCO) against 2L of an appropriate buffer. The buffer was changed after 4−5 hours and the dialysis was repeated overnight. The biotinylated protein was aliquoted, snap-frozen with liquid nitrogen, and stored at -80 °C.

**Phage Display**

An M13 phage library (Ph.D.-12, New England Biolabs) encoding random 12-mer peptides at the NH_2_ terminus of pIII coat protein (2.7 × 10^9^ sequences) was used. Biotinylated full length eIF4E was loaded onto 10 µl of steptavdin M280 magnetic Dynabeads (Invitrogen). The loaded beads were incubated with blocking buffer (20 mM HEPES pH 7.6, 0.1 M KCL, 0.5% Tween20, 2% BSA) for 1 h at room temperature, washed with buffer W (20 nM HEPES pH 7.6, 0.1 M KCL, 0.5% Tween 20), and incubated in buffer W at room temperature with 4 × 10^10^ phages. Magnetic M280 beads were the washed 8 times in buffer W. Bound phages were eluted with 0.2 M glycine (pH 2.2) and neutralized with 1 M Tris (pH 9.1). The eluted phages were amplified as instructed by the manufacturer. The selection process was repeated for three cycles. Phage plaques from the final round were picked and amplified as described by the manufacturer and sequenced.

M13 phage library (Ph.D.-C7C, New England Biolabs) encoding random 7-mer peptides flanked by two Cys was used. A 96-well microplate (Corning, #3370) was first coated with streptavidin (100 μg/mL) in 100 mM NaHCO_3_ (pH 8.4) at 4 °C overnight. After washing with 4 × 200 μL of binding buffer (50 mM Tris-HCl, 150 mM NaCl, 2 mM MgCl_2_, pH 7.4), the wells were filled with the corresponding biotinylated protein (eIF4E, eIF4A or MDM2, 20 μg/mL) in the binding buffer. After incubating at room temperature for 15 min, the microplate was washed with 4 × 200 μL of binding buffer and blocked with blocking buffer (binding buffer plus 1% BSA and 0.1% Tween-20) for 1 hour at room temperature. In parallel, the phage library was diluted to ~1.0 × 10^12^ pfu/mL in the blocking buffer. After removal of the blocking buffer from the microplate, phage library (100 μL per well) was added, incubated for 1 hour at room temperature, and washed with 4 × 200 μL of washing solution (binding buffer + 0.1% Tween-20), followed by 2 × 200 μL washing solution containing 1 mM streptavidin, and finally with 4 × 200 μL of washing solution. The bound phages were eluted for 9 min with 0.2 M glycine (pH 2.2) plus 1% BSA and neutralized with 1 M Tris (pH 9.1). The eluted phages were amplified according to manufacturer’s instruction and phage DNA was extracted by QIAprep spin Miniprep kit. The single-stranded DNA was then converted to Illumina-compatible double-stranded DNA amplicon by PCR as described previously.^3^ Sequencing was performed using the Illumina NextSeq platform (Axil Scientific). Identification of significantly enriched sequences from deep-sequencing data was performed similarly to the procedure described in Ng et al. ^4^

**Fluorescence anisotropy competition assays and K_d_ determination.**

Purified eIF4E was titrated against 50 nM carboxyfluorescein (FAM) labelled tracer peptide (Ac-KKRYSRDFLLALQK-(FAM)-NH_2_) or m^7^GTP^5-FAM^ (Jena Biosciences Cat. No: NU-824-5FM). The K_d_ (dissociation constant) for the titration of eIF4E against the tracer peptide was determined by fitting the experimental data to a 1:1 binding model equation^5,6^:

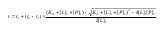

where [P] is the protein concentration, [L] is the labelled peptide concentration, r is the anisotropy measured, r_0_ is the anisotropy of the free peptide, r_b_ is the anisotropy of the eIF4E–tracer peptide complex, [L]_t_ is the total FAM labelled peptide or m^7^GTP^FAM^ concentration, and [P]_t_ is the total eIF4E concentration. The K_d_s determined for the interaction of either the tracer peptide or m^7^GTP^FAM^ with eIF4E were 50.3 nM and 149.0 nM respectively. These were used in subsequent K_d_ determinations in competition experiments to measure binding against the eIF4E cap-binding site and eIF4E:4G interface.

To determine K_d_s for compounds that disrupted the eIF4E:4G interfaces, molecules were titrated against eIF4E and the labelled peptide at set concentrations of 200 nM and 50 nM, respectively. With respect to compounds that interacted at the eIF4E cap binding site, titrations were performed with eIF4E and m^7^GTP^5-FAM^ at concentrations of 250 nM and 50 nM. Apparent K_d_ values for both competition assays were determined by fitting the experimental data to the equations shown below^35,36^:

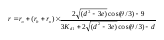

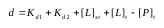

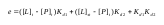

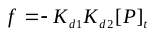

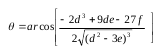

[L]_st_ and [L]_t_ denote labelled ligand and total unlabelled ligand input concentrations, respectively. K_d2_ is the dissociation constant of the interaction between the unlabelled ligand and the protein. In all competitive types of experiments, it is assumed that [P]_t_ > [L]_st_, otherwise considerable amounts of free labelled ligand would always be present and would interfere with measurements. K_d1_ is the apparent K_d_ for the labelled peptide used in the respective experiment. The tracer peptide was dissolved in DMSO at 1 mM and diluted into experimental buffer. Readings were carried out with a Envision Multi-label Reader (PerkinElmer). Experiments were carried out in PBS (2.7mM KCl, 137mM NaCl, 10mM Na_2_HPO_4_ and 2mM KH_2_PO_4_ (pH 7.4)) and 0.1% Tween 20 buffer. All titrations were carried out in triplicate. Curve-fitting was carried out using Prism 4.0 (GraphPad).

**Tryptophan quenching**

Tryptophan fluorescence quenching studies were performed using a Envision Multiplate in a black 96 well plate. Protein samples were excited at a wavelength of 290 nm and tryptophan emission was measured at a wavelength of 355 nm. Sample wells contained eIF4E at a concentration of 10 µM at a set volume of 100 ul with increasing concentrations of the relevant compounds under study. Quenching experiments were performed in PBS buffer (2.7mM KCl, 137mM NaCl, 10mM Na_2_HPO_4_ and 2mM KH_2_PO_4_ (pH 7.4)) with a final DMSO concentration of 1% (v/v).

**Construction and Assessment of VH domains with Rationally Designed Linkers**

VH-DiF clones containing different linker regions flanking the eIF4E cap-site interaction motif (EMGFF) were ordered as Ultramer double stranded oligonucleotides (IDT) containing EagI/HIndIII restriction sites. The double stranded VH-Dif encoding cassettes were then cloned into the pET22b bacteria expression vector via ligation at the EagI/HIndIII cloning sites in frame with the c-terminal polyHis affinity purification tag. VH domain constructs (R1-R3) were purified as out lined in the “**Bacterial Expression and Purification of VH-Domain constructs”** section. Purified VH clones were incubated in a 20:1 excess ratio to purified GST-eIF4E (20 µM) and incubated for 4 hours at room temperature in PBS with 1 mM DTT (see “**eIF4E expression and purification for Sortase Labelling”).** GST-eIF4E:VH domain complexes were pulled down with 20 µl of GST-beads (Thermo Fisher). Protein samples were analyzed using SDS-PAGE gel and visualized with Coomassie stain.

**Yeast Display PELE Library construction**

The pCT-CON vector was digested using SalI, NheI, and BamHI restriction enzymes (NEB) to ensure complete linearization and absence of full-length insert, thereby preventing transformation of yeast cells with parental plasmid. The PELE library of Dif-VH domains was constructed by three-step overlap extension PCR (OE-PCR). A set of 9 primers; P1_for, P2_rev to P9_rev were dissolved at 100 μM concentration and mixed in an equimolar ratio to prepare three mixed pools containing each primer at a concentration of 10 μM. The three mixed pools were denoted ‘Lib1’, ‘Lib2’, and ‘Lib3’ with each containing a primer specifically encoding a designed PELE library, P9a_for, P9b_for, or P9c for, respectively. 1 μL was taken from each mixed library and 5-fold dilution series prepared to identify the optimal primer concentration for OE-PCR. 0.4 μM of each primer was found to produce optimal yields for OE-PCR for each of the three mixed pools. The full length diF-VH domain product from each library OE-PCR reaction (Lib1, Lib2 and Lib3) was mixed in a 1:1:1 molar ratio (denoted ‘pooled PELE library’). 300 ng of the pooled PELE library and 1 μg of digested pCT-CON vector were combined with 50–100 μL of electrocompetent EBY100 yeast cells and electroporated at 0.54 kV and 25 μF using a GenePulser Xcell (Bio-Rad). Homologous recombination of the linearized vector and pooled PELE insert yielded intact plasmid. Cells were grown in YPD (1% yeast extract, 2% peptone, 2% glucose) for 1 h at 30 °C, 250 rpm. The number of total transformants was 5.7 × 10^7^ cells as determined by serial dilutions plated on SD-CAA plates (0.1 M sodium phosphate, pH 6.0, 182 g/L sorbitol, 6.7 g/L yeast nitrogen base, 5 g/L casamino acids, 20 g/L glucose). The library was propagated by selective growth in SD-CAA, pH 5.3 (0.07 M sodium citrate, pH 5.3, 6.7 g/L yeast nitrogen base, 5 g/L casamino acids, 20 g/L glucose, 0.1 g/L kanamycin, 100 kU/L penicillin, and 0.1 g/L streptomycin) at 30 °C, 250 rpm.

**Selection of Yeast Displayed PELE Library**

1 × 10^10^ cells were taken from the propagated library culture and pelleted at 2,500*g* for 5 min in 50-ml conical tubes and resuspended in SGCAA media to an absorbance of about 0.5–1 at 600 nm to induce expression of the pooled PELE library. Library induction was maintained for 24 hours at 20°C.

1 x 10^8^ induced yeast cells were pelleted at 2500g for 5 mins and the supernatant aspirated. Cells were then washed in 25 ml PBSM buffer, re-pelleted, supernatant discarded and re-suspended in 10 mls of PBSM. Biotinylated eIF4E was then added to a final concentration of 1 µM. The yeast cell suspension was then incubated at room temperature with gentle rotation on a tube rotator for 60 min, followed by 10 mins on ice.

Dif-VH domain:eIF4E complexes were then isolated using MidiMACS (Mitenyi Biotec) magnetic separation. An LS column (Mitenyi Biotec) was equilibrated with PBSM buffer at 4 °C. The yeast cell suspension was then pelleted at 2500*g* for 5 min at 4 °C, the supernatant aspirated, the yeast cells washed with 50 ml PBSM buffer and then resuspended in 5 ml PBSM buffer. 200 µl of streptavidin microbeads were added to the suspension and incubated on ice with gentle mixing for 10 minutes. Yeast cells were spun down and again washed, before being re-suspended in 50 ml PBSM buffer. Cells were then applied to the LS column in the presence of magnet. Unlabelled cells were washed from the column with 10 ml ice cold PBSM. Cells labelled with streptavidin microbeads were eluted from the column by removal from the magnetic field into a collection tube. Serial dilutions of the sorted cell suspension were plated onto SDCAA plates and incubated at 30 °C to estimate the number of cells captured by MACS. Eluted yeast cells were then propagated with addition of SDCAA (containing 100 units/ml and 100 mg/ml of Penicillin-streptomycin) media to a final volume of 500 ml and incubated overnight at 30 °C.

Magnetic sorting was then followed by 2 rounds of FACs enrichment. 1 × 10^8^ cells were taken from the propagated library culture and pelleted at 2,500*g* for 5 min in 50-ml conical tubes and resuspended in SGCAA media to an absorbance of about 0.5–1 at 600 nm to induce expression of the sorted PELE library. Cells were incubated overnight at 20°C. Induced cells were spun down at 14,000*g* for 30s, supernatant aspirated and cells washed with PBSF buffer. Yeast cells were then labelled with 500 nM of sortase labelled biotinylated eIF4E in 1 ml of PBSF buffer and incubated at room temperature for 30 mins. Cells were then pelleted by centrifugation (14,000g for 30 s at 4 °C), the supernatant aspirated and then washed with 1 ml ice-cold PBSA. Yeast were resuspended in 500 µl PBSF containing Anti-HA Ab Alexa Fluor 488 (Invitrogen, 1:100 fold dilution) and Streptavidin-phycoerythrin (ThermoFisher Scientific, 1:100 fold dilution) and incubated for 30 mins. Cells were then pelleted at 14,000*g* for 30 s at 4 °C, washed with 1 ml PBSF buffer and resuspended in 2.0 mL PBSF. Cells positive for anti-HA and eIF4E were selected and sorted using an Aria (Becton Dickinson) cytometer. Collected cells were propagated in SDCAA at 30 °C and a second round of FACs selection performed after yeast induction with 1 × 10^8^ cells as described.

After the final round of FACs selection, serial dilutions of the sorted cell suspension were plated onto SDCAA plates and incubated at 30 °C until the appearance of yeast colonies. X colonies of yeast were individually picked and then propagated in X mls of SCDAA. Plasmid DNA was then isolated using the Zymoprep kit II (following the manufacturer’s instructions), cleaned using the Qiagen PCR Purification kit, and transformed into DH5α (Invitrogen) cells. Purified plasmids were then sequenced using BigDye chemistry.

**Assessment of Enriched Yeast PELE Library for Specific Binders to the eIF4E Cap-Binding Site**

Before the second round of FACs sorting an additional subset of 5 × 10^7^ cells was induced in 5 ml of SGCAA media. Yeast were then pelleted, washed in 1 mL PBSF resuspended in PBSF to a density of 1 x10^7^ cell per ml. 1 ml of yeast suspension was added to three individual tubes. Purified sortase biotinylated eIF4E was then added to each sample at a concentration of 2 µM and samples were incubated at 20 °C for 1 hour. Purified sortase biotinylated eIF4E was then added to each sample at a concentration of 0.2 µM either in combination with 50 µM m^7^GTP (Sigma-Aldrich), 50 µM of purified 4E-BP1^4ALA^ or 50 µM of VH-1C5^M4^, followed by sample incubation at 20 °C for 1 hour. 4E-BP1^4ALA^ and VH-1C5^M4^ were purified as described previously.^7^ Cells were then pelleted by centrifugation (14,000g for 30 s at 4 °C), the supernatant aspirated and then washed with 1 ml ice-cold PBSA. Yeast were resuspended in 500 µl PBSF containing Anti-HA Ab Alexa Fluor 488 (Invitrogen) and Streptavidin-phycoerythrin (ThermoFisher Scientific) and incubated for 30 mins. Cells were then pelleted at 14,000g for 30 s at 4°C, aspirate supernatant and wash with 1 ml PBSF buffer. Each sample was then analysed by flow cytometry using Aria (Becton Dickinson) cytometer.

**VH Domain Expression and Purification Assessment**

VH domains were amplified and cloned into the pET-22b (+) vector (Novagen) using the in-Fusion cloning method (Takara Bio) as described earlier. These VH domains were expressed into E.coli BL21 (DE3) cells. Cells were grown at 37 °C and induced protein expression overnight at 25 °C by 0.5mM Isopropyl-β-D-thiogalactoside (IPTG). For assessment of clones, cells from 20 ml cultures was harvested, and lysed by sonication in lysis buffer (25mM HEPES pH 7.5, 300mM NaCl, 20mM imidazole, 1mM DTT) supplemented with protease inhibitor cocktail. After centrifugation, the supernatant containing soluble proteins was loaded into Ni-NTA spin column (Qiagen). The column was washed twice using lysis buffer and then eluted with 25mM HEPES at pH 7.5, 300mM NaCl, 1mM DTT and 500mM imidazole. To assess protein solubility of different VH domains, the eluted proteins were analyzed by SDS-PAGE gel and stained with coomassie blue.

**Bacterial Expression and Purification of VH-Domain constructs**

VH-Dif sequences were ordered as gene fragments from Integrated DNA Technologies (IDT). Coding sequences were PCR amplified and cloned directly into the bacterial expression vector pET-22b(+) with an in-frame C-terminal six-hisitidine tag using thr BamHI/XhoI cloning site. Each VH-domain plasmid was separately transformed into *E. coli* BL21 (DE3) cells and used to inoculate 10 ml of LB broth (containing 100µg/ml) started culture, which were incubated overnight before being used to seed 1000 ml of fresh LB broth. Bacterial cultures were grown at 37 °C and when they reached a OD_600_ of 0.6–0.8, the cells were induced with a final concentration 0.5 mM IPTG and incubated overnight at 25 °C. Cells were harvested by centrifugation at 5,000 rpm for 10 minutes and pellets were resuspended in lysis buffer (25 mM HEPES pH 7.5, 300 mM NaCl, 20 mM imidazole, 1 mM DTT) and then sonicated for 5 minutes. Bacterial supernatants were then filtered through 1.2 μm syringe filters. Proteins were purified through a standard two-step protocol: first, supernatant was loaded onto a pre-equilibrated 1 ml HisTrap column (Cytiva Lifesciences), which was then extensively washed with buffer A (25 mM HEPES pH 7.5, 300 mM NaCl, 1mM DTT) and then eluted with an imidazole gradient; second, the eluted proteins were subjected to gel filtration chromatography on a Superdex 75 column (Cytiva Lifesciences) using PBS buffer containing 1 mM DTT. Protein fractions were analysed by SDS page gel and concentrated. Protein concentration was determined using absorbance at A280 nm.

**Protein Crystallization**

The eIF4E:EE-02 and eIF4E:V_H_-Dif^CAP^-01 complexes were crystallized by vapour diffusion using the hanging drop method. For crystallization, the eIF4E:EE-02 complex was prepared by direct addition of a 100mM DMSO stock solution of EE-02 to purified eIF4E recombinant protein (dialysed in 10 mM HEPES 7.6, 100 mM KCL buffer) to generate a final solution of 200 µM eIF4E and 300 µM EE-02 with a residual DMSO concentration of 0.3% (v/v). The sample was then spun down using a tabletop centrifuge at 13,000g after overnight incubation at 4 °C and the supernatant used for crystallization. The eIF4E: V_H_-Dif^CAP^-01 complex solution for crystallization was prepared by dialysing both proteins into 10 mM HEPES 7.6, 100mM KCL and 1mM DTT buffer and mixing them to give final respective concentrations of 100 µM and 200 µM. Hanging drops were set-up in a pre-greased VDX48 plate (Hampton, USA) with 1 µl of the respective crystallization sample mixed with 1 µl of the mother-well solution. eIF4E:EE-02 crystals grew over a period of one week in 0.2 M Potassium chloride, 20% (w/v) PEG 3350. eIF4E:V_H_-Dif^CAP^-01 crystals grew over a similar period of time but in 0.1 M TRIS.HCl pH 8.5, 25% (v/v) PEG 550 MME. For X-ray data collection at 100 K, crystals for both sets of crystallization conditions were transferred to an equivalent mother liquor solution containing 25% (v/v) glycerol and then flash frozen in liquid nitrogen.

**Data collection and refinement**

X-ray diffraction data was collected at the Australian synchrotron (MX1 beamline) using a CCD detector, and integrated and scaled using XDS. The initial phases of the EE02 complexed crystal of eIF4E were solved by molecular replacement with the program PHASER^8^ using the human eIF4E structure (PDB accession code**:**  4BEA) as a search model. With respect to the eIF4E:V_H_-Dif^CAP^-01 co-crystal the VH domain structure (PDB accession code**:**  5TDP, chain B) was also included in the PHASER molecular replacement search as an independent search model. The starting models were subjected to rigid body refinement and followed by iterative cycles of manual model building in Coot and restrained refinement in Refmac 6.0.^9^ Models were validated using PROCHECK^10^ and the MOLPROBITY webserver.^11^ Final models were analysed using PYMOL (Schrödinger). See table S1 for data collection and refinement statistics. The eIF4E complex structures with EE-02 and eIF4E V_H_-Dif^CAP^-01 have been deposited in the PDB under the submission codes 7EZW and 7F07, respectively.

**Isothermal Titration calorimetry (ITC)**

ITC measurements were performed with the Affinity ITC (TA Instruments, USA) at 25 °C. The purified proteins were buffer exchanged into 1×PBS, pH 7.2 with 0.001% Tween-20 using 7K MWCO Zeba spin desalting column (ThermoFisher scientific). 10-30 μM of eIF4E protein was loaded into the sample cell, and 100-300 μM of VH domains were titrated into eIF4E protein, over 15-20 injections of 2.5 μL. All experiments were conducted in duplicate. Calorimetric data were analysed with NanoAnalyze software using a one-site binding model. Correction for the enthalpy of ligand dilution was carried out by subtracting a linear fit from the last three data points of the titration, after the interaction had reached saturation.

**Molecular Dynamics Simulations**

The X-ray resolved VH-DIF^CAP^-01: eIF4E and EE-2: eIF4E complex state structures, along with the free VH-DIF^CAP^-01 domain and EE-2 cyclic peptide derived from the respective complexes were subjected to molecular dynamics simulations in AMBER 18^12^ using all-atom ff14SB^13^ force field parameters. The N-termini of eIF4E and VH-DIF^CAP^-01 were capped with the ACE functional group, while the C-termini of VH-DIF^CAP^-01 and EE-2 were capped with NME and NHE functional groups respectively. The disulphide bond between residues C2 and C10 in the EE-2 peptide was maintained using the “bond” command in the tleap module of AMBER 18. All the water molecules resolved in the crystal structures were retained for the simulations. The four systems (VH-DIF^CAP^-01: eIF4E, EE-2: eIF4E, free VH-DIF^CAP^-01 and free EE-2) were placed inside a truncated octahedral box and solvated with TIP3P^14^ water by setting a minimum distance of at-least 8 Å between any solute atom and the edge of the box. The electroneutrality of the respective systems was achieved by adding appropriate number of counterions. These systems were then energy minimized using steepest descent and conjugate gradient algorithms, heated to a temperature of 300 K in the NVT ensemble and equilibrated for 500 ps in the NPT ensemble with 1 atm pressure. Production dynamics for VH-DIF^CAP^-01: eIF4E and EE-2: eIF4E complexes were carried out in triplicates for 200 ns each (cumulative simulation time of 1.2 μs) starting with different initial velocities, while that for free VH-DIF^CAP^-01 domain and EE-2 cyclic peptide was run for 1 μs each. All the simulations in the production stage were carried out under NPT conditions. Electrostatic calculations, regulation of temperature and pressure along with the constraining of bonds to hydrogen atoms during the simulations were employed as previously described by Lama et al.^15^

**Analysis of binding energy and water occupancy during the molecular dynamics simulations**

Residue-wise decomposition analysis was carried out using the MM/GBSA (Molecular Mechanics / Generalized Born Surface Area)^16^ scheme through the MMPBSA.py script available in AMBER 18. The 3 simulated trajectories of each complex, ie VH-DIF^CAP^-01: eIF4E and EE-2: eIF4E, were concatenated (200 ns per trajectory) into a composite trajectory of 600 ns and from this, 1000 snapshots at equal intervals, were extracted. Water molecules and counterions were removed from these structures and solvation effects were estimated using the implicit Generalized Born Solvation Model (IGB=2) with salt concentration set to 150 mM. The water occupancy map for VH-DIF^CAP^-01: eIF4E and free VH-DIF^CAP^-01 simulations was generated using the “grid” command in the CPPTRAJ module of AMBER 18 with cubic grid cells of size 0.5 Å. The water density within each grid cell was computed and plotted using the volume viewer menu as an isosurface representation in the UCSF Chimera visualization software.^17^

**Cell Biology**

***Plasmid and Reagents***

All plasmids were purchased from Addgene unless indicated otherwise. Mutant VH domains were generated with In-fusion mutagenesis kit (Clontech) and then cloned into either a pCDNA3.1 vector (Thermo Fisher Scientific) harbouring a C-terminal 3× FLAG tag via NheI/BamHI sites or into NanoBit plasmids using the NanoBit PPI starter system (Promega, see manufacturer instruction) to allow mammalian cell expression studies. For lentivirus production, the VH-3xFLAG cassette was sub-cloned into the lentiviral expression vector pCW57 via BamHI/AvrII cloning sites. pcDNA3-rLuc-polIRES-fLuc (bicistronic reporter)^ref^, eIF4E and eIF4G^604–646^ NanoBIT and 4EBP1^4ALA^ mutant plasmid generation has been described previously in the literature.^18^

***Cell Culturing Conditions***

All cell lines were cultured in DMEM cell media supplemented with 10% foetal calf serum (FBS) and penicillin/streptomycin. Mammalian cells were maintained in a 37 °C humidified incubator with 5% CO_2_ atmosphere.

***Immunoprecipitation and m^7^GTP pull down experiments***

Twenty-four hours prior to transfection or drugging, cells were seeded at a cell density of 1000,000 (HEK293) cells per well of a six-well plate (ThermoFisher Scientific). Transfections were performed using Lipofectamine 3000 (ThermoFisher Scientific) with either 2 μg or the indicated amount of plasmid vectors per well according to the manufacturer’s instructions. After a 48-hours (or as indicated in the relevant figure) incubation period, the cell media was then removed and the cells washed with PBS saline. Cells were directly lysed in the wells with 300 μl of lysis buffer containing 20 mM Hepes pH 7.4, 100 mM NaCl, 5 mM MgCl_2_, 0.5% NP-40, 1 mM DTT, with protease (Roche) and phosphatase (Sigma-Aldrich) inhibitor cocktail sets added as outlined by the manufacturer’s protocols. Cellular debris was removed by centrifugation, and the protein concentration was then determined using the BCA system (Pierce). m^7^GTP pulldown and FLAG immunoprecipitation experiments were performed with 200 μg of cell lysate, which was either incubated with 20 μl of m^7^GTP (Jena Bioscience) or anti-FLAG M2 antibody (Roche) immobilised agarose beads for 2–4 hours at 4 °C on a rotator. Beads were then washed four times with lysis buffer containing no protease or phosphatase inhibitors. This was then followed by the addition of Laemlee buffer (2×) and the beads boiled for 5 min at 95 °C. Samples were centrifuged and the supernatant removed for western blot analysis.

***NanoBit*** ***complementation Assay***

For NanoBIT (PROMEGA) system development and validation, opaque 96-well plates were seeded with 30,000 HEK293 cells per well in DMEM and 10% FCS and transfected with 30 ng total DNA of the two NanoBit plasmid vectors and 100 µg of the indicated plasmid per well using FUGENE6 (Roche). 48 hours after transfection, the medium was replaced with 100 μl of Opti-MEM cell media containing 0% FCS with no added red phenol (Thermo Fisher Scientific). To screen indicated compounds using the NanoBIT system in live and permeabilized cells, 6 well plate were seeded with 1300,000 cells per well in DMEM and 10% FCS and transfected with 2 ug total DNA of the two NanoBit plasmid per well. After 24 hrs, transfected cells were trypsinised and re-suspended in Opti-MEM media with 10% FCS. Cells were then spun down at 1000 rpm for 5 minutes at room temperature. Supernatant was then discarded and cells re-suspended to a density of to 220,000 cells per ml in Opti-MEM I reduced serum containing 10% FCS with no added red phenol. 100 μl of cells were added to the wells of a white opaque 96-well plate and incubated for 24 hours at 37 ̊C, 5% CO2. For assessment of permeabilised or live cells, cell medium was replaced with 90 ul of serum free Opti-MEM media that either contained or did not contain 50 ug/ml digitonin, respectively. Live or permeabilized cells were then treated with either 10 μl of a 10% v/v DMSO vehicle control in FPLC grade water or a suitable 2-fold dilution series of the compound under study in a 10-fold higher stock concentration (containing 10% DMSO and FPLC grade water solution). 96 well were then incubated for 3 hrs at 37 ̊C, 5% CO_2_. Luminescence activity was assayed as described elsewhere^18^ using an Envision Multi-Plate reader.

***Cap-Dependent Translation Assay***

Opaque 96-well plates were seeded with 30,000 HEK293 cells per well in DMEM and 10% FCS. Transfections were performed using FUGENE6 (Roche) with 30 ng of the bicistronic reporter (pcDNA3-rLuc-polIRES-fLuc) plasmid and 75 or 150 ng of the indicated plasmid. 48 hours after transfection, Renilla and firefly luminescence activity was determined using the Dual Glo Luciferase Assay System (PROMEGA). Luminescence readings were performed using an Envision Multi-plate reader (PerkinElmer).

***Protein Expression Analysis***

Transfected HEK293 cells (prepared as described in the NanoBit and Cap-dependent translation Experiments sections) were seeded with 30,000 cells per well in 96-well plates. After an incubation period indicated in the relevant figure, cells were washed with PBS and directly lysed in the wells of the plate with 50 μl of cell lysis buffer (20 mM Hepes pH 7.4, 100 mM NaCl, 5 mM MgCl_2_, 0.5% NP-40, 1 mM dithiothreitol) containing the protease (Roche) and phosphatase (Sigma-Aldrich) inhibitor cocktail sets (added as outlined by the manufacturer’s protocols). Cellular debris was separated by centrifugation. Samples were analysed by western blot without further quantification.

***Western Blot analysis***

Samples were resolved on midi or mini Tris-Glycine 4–20% gradient gels (Bio-Rad) according to the manufacturer’s protocol. Western transfer was performed with an Immuno-blot PVDF or nitrocellulose membrane (Bio-Rad) using a Trans-Blot Turbo system (Bio-Rad). Western blots were then performed. Antibodies against peIF4E^S209^,4EBP1, cyclin D1 and FLAG were purchased from Abcam or Sigma, respectively. All other antibodies used were purchased from Cell Signalling Technology. Β-actin levels were measured to ensure equal loading.

***Cell Proliferation Assay***

A375 cell lines were plated in 96-well clear bottom plates at a density of 4000 cells per well in 200 ul DMEM and 10% FCS medium. After 24 hours, cell media was replaced with 200 µl of medium containing doxycycline at 1 ug/ml. Cell confluence and cell growth was then measured continuously over 7 days using an IncuCyte FLR instrument (EssenBioscience).
